# A novel product-tensor decomposition of two independent datasets can reveal a shared brain–skin transcriptional program under restraint stress

**DOI:** 10.64898/2026.11.21.768350

**Authors:** Y-H Taguchi, Turki Turki

**Affiliations:** Department of Physics, Chuo University, 1-13-27 Kasuga, Bunkyo-ku, Tokyo 112-8551, Japan. Bioinformatics Center, Institute for Chemical Research, Kyoto University, Kyoto, 611-0011, Japan.; Department of Computer Science, King Abdulaziz University, Jeddah 21589, Saudi Arabia.

**Keywords:** tensor decomposition, unsupervised feature extraction, brain–skin axis, restraint stress, multi-omics integration, higher-order singular value decomposition

## Abstract

**Background:** Psychological stress acts on the brain and on peripheral organs at once, yet the two are rarely measured in the same animals. Datasets from separate studies then share a gene list and nothing else: no animals, no batch, no sequencing run. Methods that integrate such datasets assume a correspondence between samples that does not exist, while intersecting two lists of differentially expressed genes abandons the requirement that a gene respond in both tissues at once.

**Results:** We introduce a product-tensor formulation of tensor-decomposition-based unsupervised feature extraction. The outer product of the two profiles along their shared feature axis pairs every sample of one study with every sample of the other, so that no matched samples, no batch correction and no differential expression step are required, and a gene is selected only if it carries a contrast on both sides. Applied to mouse ventral hippocampus and mouse dorsal skin under restraint stress, two studies sharing only the stressor, the method recovers 977 to 1223 genes at each of six sampling times, with 83 to 87 percent of the squared length of the gene-mode singular vector lying outside the fitted null. The selected genes are attributed to both tissues by independent reference atlases and are enriched for extracellular matrix organisation, epithelial–mesenchymal transition and TNF/NF-***κ***B signalling. Their relative sign separates a concordant leukocyte programme from a discordant mesenchymal one. Five of the six times replicate in a cohort stressed for twice as long, at 5.0 to 6.0 times chance, and adding chromatin accessibility as a further mode leaves the selection unchanged.

**Conclusions:** Two tissues profiled by different laboratories under one stimulus can be integrated without inventing a correspondence between their samples. On this pair of datasets the construction recovers a brain–skin transcriptional response to restraint stress that separates by direction into an immune and a mesenchymal arm, and that replicates in animals the method never saw.

## 1 Background

The response to psychological stress is not confined to the brain. Glucocorticoids and catecholamines released by the hypothalamic–pituitary–adrenal axis and by the sympathetic nervous system reach every tissue, and the skin in particular possesses a local neuroendocrine apparatus of its own [1–3]. The clinical consequences are familiar: stress precedes flares of psoriasis and atopic dermatitis and increases susceptibility to cutaneous infection. Recently, Chan et al. [4] established a causal chain for the latter, showing that restraint stress acts through adrenergic signalling and TGF-*β* to suppress cathelicidin production by dermal fibroblasts, and describing the result explicitly as a brain–skin axis.

Evidence of this kind is necessarily built one arm at a time. A study that dissects the cutaneous arm does not profile the brain, and a study that charts the hippocampal transcriptional response to the same stressor [5] does not profile the skin. The two datasets therefore share a stimulus and a gene list, but not a single animal, a single batch or a single sequencing run. Asking whether the two tissues engage a common transcriptional program is natural, but the usual tools do not answer it. Differential expression followed by overlap of the two gene lists discards everything except membership and is dominated by the arbitrary thresholds applied on each side. Correlating fold changes requires that the contrasts be commensurable, which they are not when the designs differ in replicate number, platform and normalisation. Methods that integrate omics layers — canonical correlation, joint matrix factorisation, multi-omics factor analysis — assume samples in common, which is precisely what is missing here.

Tensor-decomposition-based unsupervised feature extraction (TD-based UFE) [6, 7] selects features without reference to a class label, by forming a tensor from the data, decomposing it with the higher-order singular value decomposition (HOSVD) [8], identifying the components whose sample-mode singular vectors carry the contrast of interest, and testing the corresponding feature-mode singular vector against a Gaussian null. We have applied it to hippocampus and colon in cognitive ageing (Taguchi, Turki and Akutsu, manuscript in preparation), but there the two tissues came from one study.

Here we extend TD-based UFE to the case where nothing but the feature axis is shared. Let *x_ijk_ ∈* R*^N×^*^2*×K*^ be the measured expression of gene *i* in the brain dataset, where *i* = 1*, …, N* runs over the *N* genes common to the two platforms, *j ∈ {*1, 2*}* labels the two experimental conditions (control, stressed) and *k* = 1*, …, K* labels the biological replicates. Let *y_isd_ ∈* R*^N×^*^2*×D*^ be the expression of the same gene *i* in the skin dataset, with *s ∈ {*1, 2*}* its two conditions and *d* = 1*, …, D* its replicates. The two datasets share the index *i* and nothing else: *k* and *d* run over different animals in different laboratories, and *K* and *D* need not be equal. We build the *product tensor*

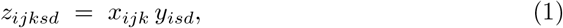

whose four sample modes are the two designs taken separately and whose single feature mode, *i*, is the gene list they share. No sample correspondence is assumed or required, because Eq. (1) pairs every brain sample (*j, k*) with every skin sample (*s, d*). A gene obtains a large loading only if it carries a contrast on *both* sides; the product is multiplicative, so a strong response in one tissue combined with none in the other cancels. Figure 1 shows the construction.

**Fig. 1.**
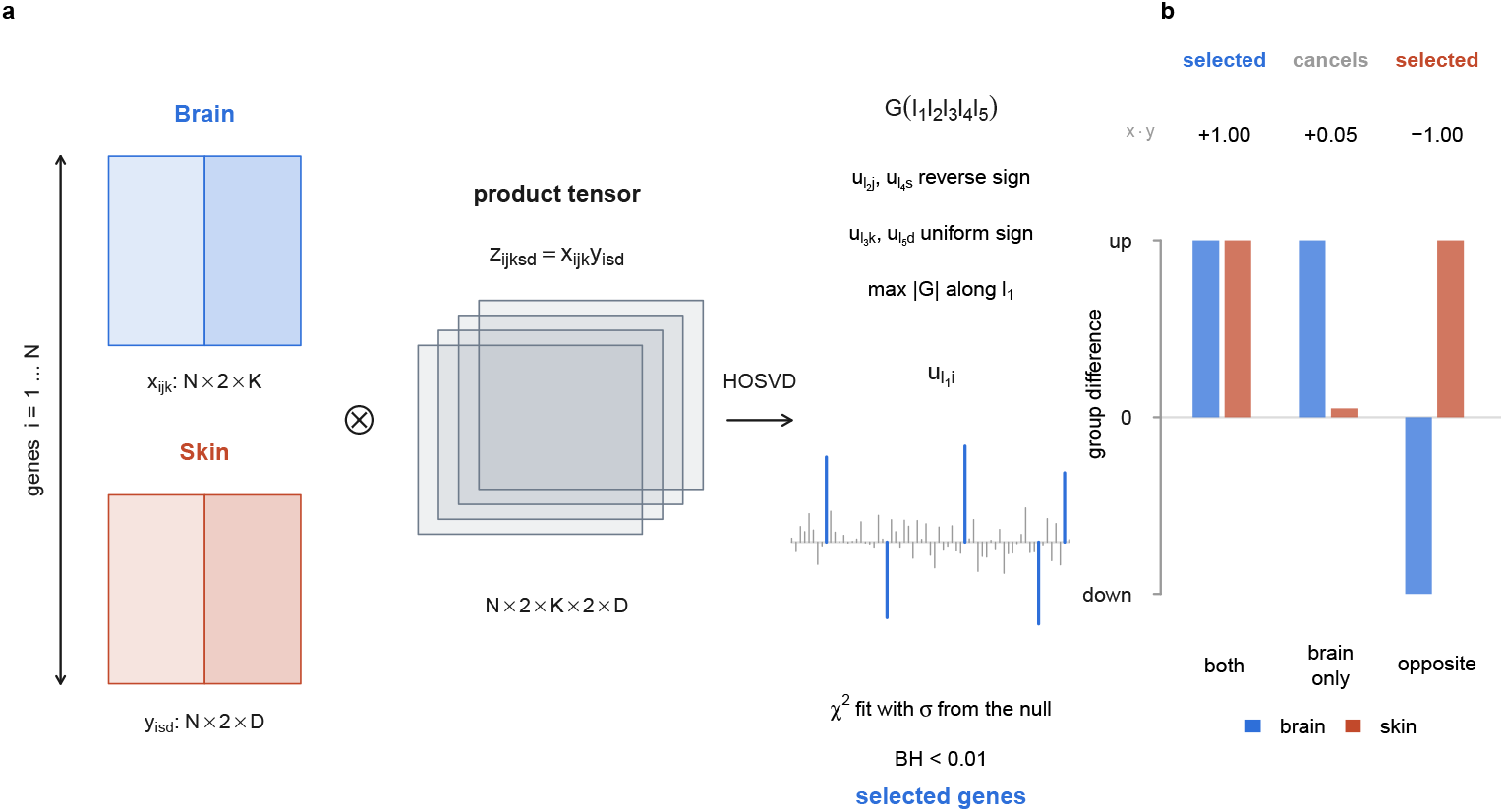
The product tensor. (**a**) The brain array *x_ijk_* and the skin array *y_isd_* share only the gene index *i*; their outer product along that index gives a five-mode tensor in which every brain sample (*j, k*) is paired with every skin sample (*s, d*). HOSVD returns the core *G*(*ℓ*_1_*ℓ*_2_*ℓ*_3_*ℓ*_4_*ℓ*_5_) and the singular vectors; the component is chosen by the sign conditions on the sample modes and by max *|G|* along *ℓ*_1_, and its gene-mode vector *u_ℓ_*_1_ *_i_* is tested against a Gaussian whose width is fitted from the null region. (**b**) Why the product selects. A gene with a group difference in both tissues gives a large product and is selected; a gene that responds in the brain alone gives a product near zero and is not; a gene that moves in opposite directions is selected with the opposite sign, which is what the direction split of Fig. 5 reads.

We apply this construction to the ventral hippocampus of mice subjected to restraint stress [5] and to the dorsal skin of mice subjected to restraint stress and infected with methicillin-resistant *S. aureus* [4]. We report that a coherent gene program is recovered at all six hippocampal sampling times, that it is attributed to both tissues by independent reference atlases, that it separates into a same-direction inflammatory arm and an opposite-direction mesenchymal arm, and that it replicates in an independent cohort of animals stressed for twice as long.

## 2 Methods

### 2.1 Data

Hippocampal data were obtained from GEO accession GSE292224 (bulk RNA-seq; part of SuperSeries GSE292240) [5]: 19 432 genes *×* 190 samples of mouse ventral hippocampus. Animals received 1 h 30 of restraint in a 50 mL tube, preceded by 10 or 20 days of daily restraint (CRS) or by handling only; tissue was taken at 0, 45 min, 1 h 30, 3 h, 5 h 30 and 24 h from the onset of the final restraint. Cutaneous data were obtained from GEO accession GSE270713 [4]: peri-lesional dorsal skin of mice infected with MRSA on day 0 and sacrificed on day 2, with restraint stress of 3 h/day for 3 days. Only sham-operated animals were used (Sham S A 1/2/3 versus Sham S A Stress 1/2/3); the adrenalectomised samples of the same series were not used. Chromatin-accessibility data were obtained from GSE292231 (bulk ATAC-seq, same SuperSeries), which covers 0 min and 45 min only, and whose animals are identical to those of the RNA-seq, the contralateral hemisphere having been used.

### 2.2 Preprocessing

Counts were transformed by log_2_(*x* + 1) and each column (sample) was normalised to unit *ℓ*_2_ norm. No per-gene scaling and no centring were applied. 14 689 genes are common to the two platforms; four Y-linked genes (*Ddx3y*, *Eif2s3y*, *Uty*, *Kdm5d*) were removed, leaving *N* = 14 685. The justification for removal is that all cutaneous animals are male, so *Ddx3y* varies only between 77 and 101 across the twelve skin samples and carries no cutaneous contrast; before removal these four genes occupied ranks 1, 2, 6 and 7 in *|u_ℓ_*_1_ *_i_|* at 45 min. *Xist* is absent from the common gene set, so sex was called from the Y-linked genes alone: their summed expression is strongly bimodal (8–25 versus 1600–3000), and every group is balanced 4 male and 4 female (3 versus 4 at 5 h 30).

### 2.3 Product tensor and decomposition

With *x_ijk_* the brain profile of gene *i ∈ {*1*, …, N}* in condition *j ∈ {*1, 2*}* and replicate *k ∈ {*1*, …, K}*, and *y_isd_* the skin profile in condition *s ∈ {*1, 2*}* and replicate *d ∈ {*1*, …, D}*, the product tensor *z_ijksd_* is defined by Eq. (1), of shape *N ×* 2 *× K ×* 2 *× D* with *N* = 14 685, *K* = 8 (*K* = 7 at 5 h 30) and *D* = 3. Replicate pairs (ctl*_k_,* trt*_k_*) on the brain side were formed within sex. HOSVD was computed with rTensor::hosvd [8, 9] at ranks (20, 2*, K,* 2*, D*), giving the core tensor *G*(*ℓ*_1_*ℓ*_2_*ℓ*_3_*ℓ*_4_*ℓ*_5_) and the singular vectors *u_ℓ_*_1_ *_i_*, *u_ℓ_*_2_ *_j_*, *u_ℓ_*_3_ *_k_*, *u_ℓ_*_4_ *_s_*, *u_ℓ_*_5_ *_d_* of Eq. (4); modes 2 to 5 are at full rank, so *u_ℓ_*_1_ *_i_* and *G* for *ℓ*_1_ = 1*, …,* 20 are unchanged by the choice of gene rank (the decompositions are nested).

A component was admissible if *u_ℓ_*_2_ *_j_* and *u_ℓ_*_4_ *_s_* (the two condition modes) change sign between levels and *u_ℓ_*_3_ *_k_* and *u_ℓ_*_5_ *_d_* (the two replicate modes) are of uniform sign. Among admissible components, the core tensor was consulted *along ℓ*_1_ *only* and the largest *|G|* taken. Components sitting exactly at the rank boundary (*ℓ*_1_ = 20) were excluded as artefacts of truncation: at 5 h 30 the apparent maximum was at *ℓ*_1_ = 20, and a full-rank decomposition (84 components) showed the true maxima to be at *ℓ*_1_ = 23 and *ℓ*_1_ = 21; we retained rank 20 and used the largest interior component, *ℓ*_1_ = 7. No other time point was affected.

### 2.4 Feature selection

The null model is that a gene not contributing to the component has a loading drawn from a zero-mean Gaussian of unknown width *σ*. With *ℓ*_1_ the selected gene component, *P* -values were therefore assigned by

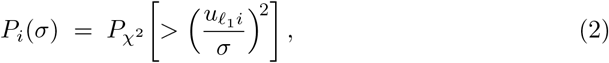

*P_χ_*_2_ being the upper tail of the *χ*^2^ distribution with one degree of freedom.

The width *σ* is not known in advance and is fitted from the data, as follows. If *σ* were correct, the *P_i_* of the genes that obey the null would be uniformly distributed on (0, 1), so their histogram would be flat. Genes that do not obey the null concentrate near *P_i_* = 0 and would spoil such a histogram; we therefore restrict attention to *P_i_ >* 0.01, which the signal does not reach, and measure flatness there. Explicitly, for a given *σ* we compute all *P_i_* from Eq. (2), take those with *P_i_ >* 0.01, bin them into 100 equal-width bins, and evaluate the standard deviation of the 100 bin counts,

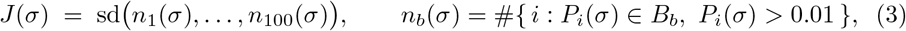

where *B_b_* is the *b*-th bin. A flat histogram makes every *n_b_* equal and *J* minimal, so *σ*^ = arg min*_σ_ J* (*σ*). *J* was evaluated on a logarithmic grid of 200 values of *σ* spanning 0.05 to 5 times sd(*u_ℓ_*_1_ *_i_*); the grid minimum was taken, the local maximum immediately to its left located, and optimize run on the interval between that maximum and the upper end of the grid. *J* rises steeply on both sides of the minimum (Fig. 2a), which is therefore well determined on the scale that matters. The bin counts are integers, however, so *J* is a step function, and on a scale of parts per million in *σ*it carries many shallow local minima; the grid spacing and the default tolerance of optimize smooth over them, and both are part of the specification rather than approximations to be refined. The residue is that a difference in *u_ℓ_*_1_ *_i_* below printing precision — a different BLAS, say — can move *σ*^ by a few parts per million, which moves genes whose adjusted *P* lies within about 10*^−^*^5^ of the threshold across it. Re-running the published pipeline in a second implementation reproduced eleven of the twelve runs of Tables 1 and 3 exactly; at 1 h 30 it returned 1196 genes rather than 1194, the two additional genes having adjusted *P* of 0.009995 and 0.009999. Gene counts should therefore be read as carrying an uncertainty of a few genes, and the released code checks them within that tolerance. No part of the procedure uses the group labels.

**Fig. 2.**
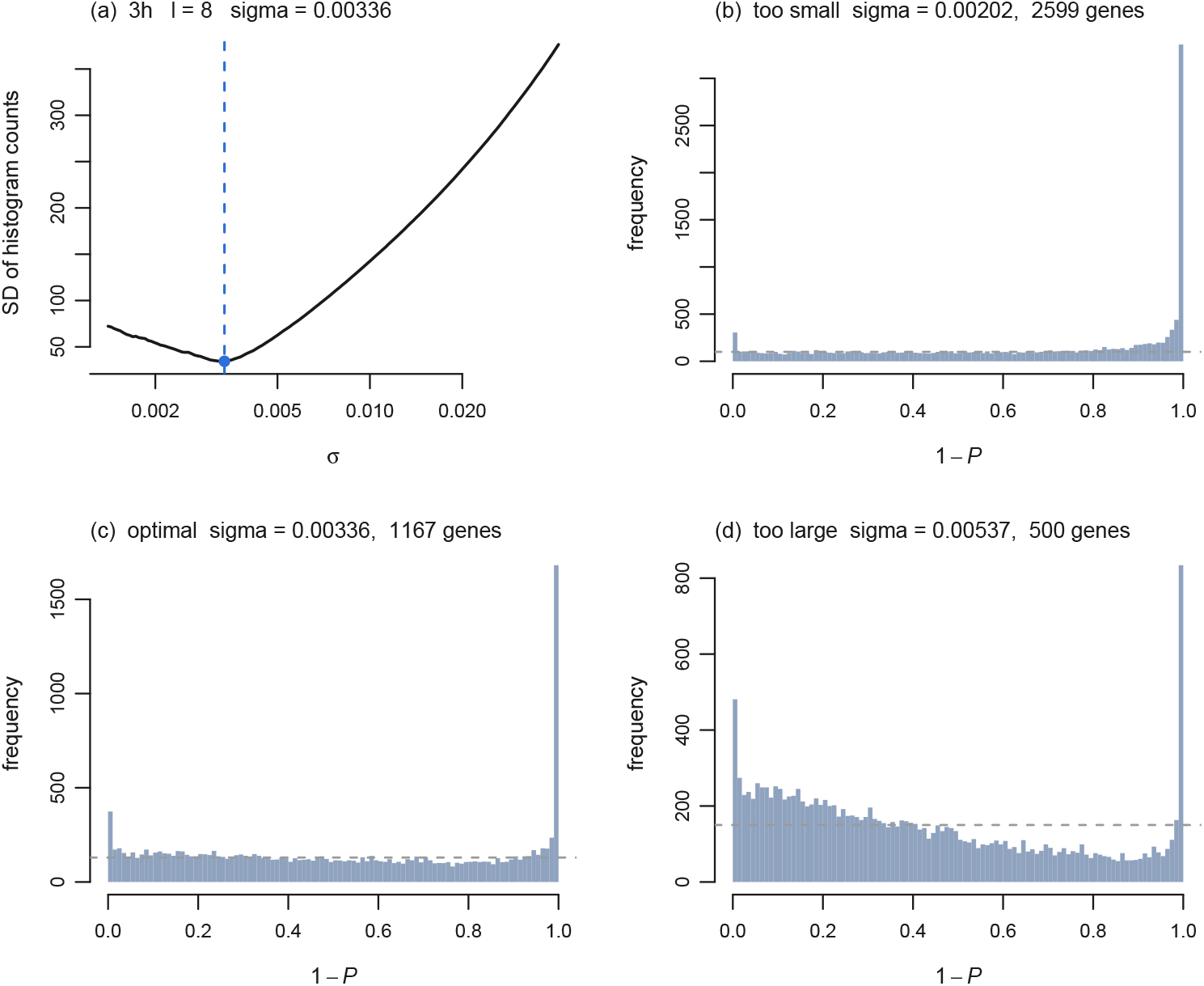
Optimisation of the null width σ at 3 h. **(a)** Objective function *J* (*σ*) of Eq. (3) against *σ* on a logarithmic axis. **(b–d)** Histograms of 1 *− P* at 0.6*σ*, *σ* and 1.6*σ*; the dashed line is the mean count over the null region. At the optimum the null region is flat, while the excess near 1*−P* = 1 is retained. The peak at 1 *− P* = 0, which differential-expression pipelines usually remove, is not removed here.

**Fig. 3.**
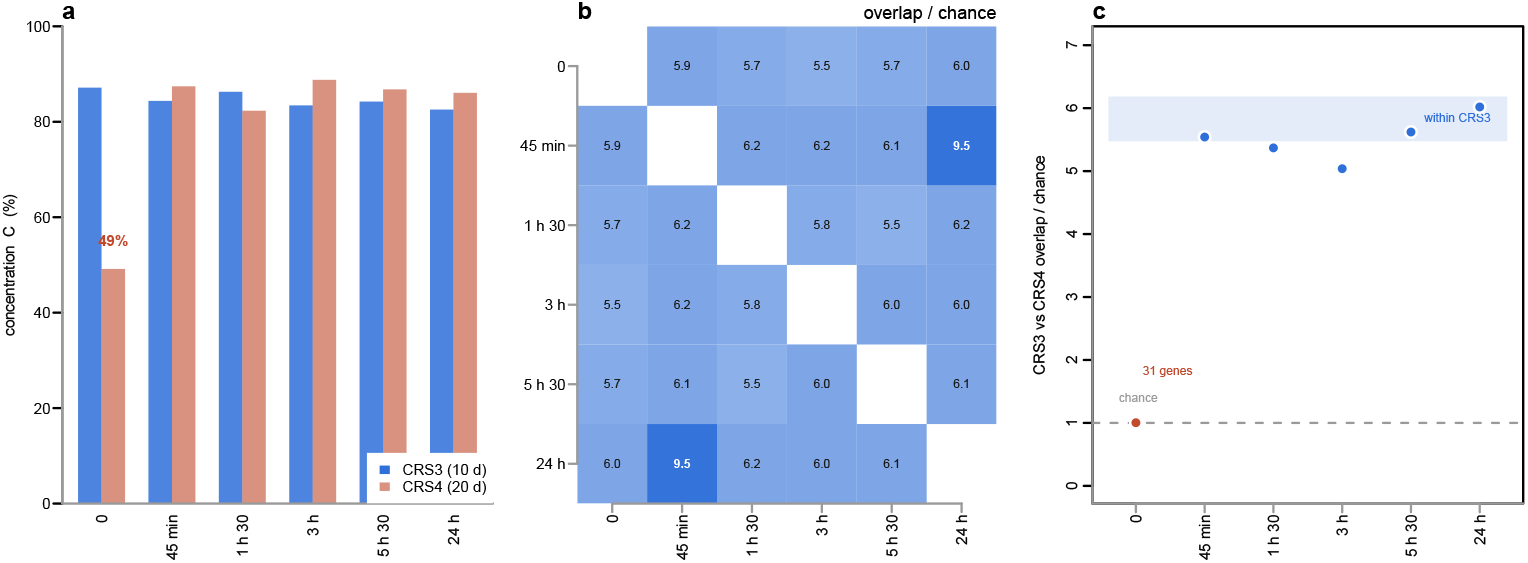
The same program is recovered at every time point and in an independent cohort. (**a**) The concentration *C* of Eq. (6) for the 10-day cohort (CRS3) and the 20-day cohort (CRS4). Eleven of the twelve runs lie between 82% and 89%; the exception is CRS4 at 0 min. (**b**) Pairwise overlap between the six CRS3 gene sets, as a multiple of the chance expectation *n_a_n_b_/N*. Fourteen of the fifteen pairs lie between 5.48 and 6.18. The exception, 45 min against 24 h, is the one pair that selects the same component. (**c**) Overlap between CRS3 and CRS4 at the same time point, against the same chance expectation. The shaded band is the range of panel **b**: matching the time point across cohorts is no more concordant than changing the time point within one. At 0 min the overlap is 31 genes, exactly the chance level.

**Table 1.** Component selection and gene counts at the six hippocampal sampling times (CRS3 cohort, 10 days of chronic restraint). *K*, number of replicate pairs on the brain side; *ℓ*_1_, index of the selected gene component; *G*, the corresponding core-tensor element; *C*, concentration of the gene-mode vector, Eq. (6). “Unique” counts genes selected at that time point and at no other.

| Time point | *K* | ℓ<sub>1</sub> | *G* | *σ* | *C* | BH < 0.01 | Unique |
|---|---|---|---|---|---|---|---|
| 0 min | 8 | 13 | +3.099 × 10<sup>−6</sup> | 0.002958 | 87.1% | 1223 | 251 |
| 45 min | 8 | 5 | −4.811 × 10<sup>−6</sup> | 0.003262 | 84.4% | 999 | 84 |
| 1 h 30 | 8 | 8 | +4.454 × 10<sup>−6</sup> | 0.003056 | 86.3% | 1194 | 223 |
| 3 h | 8 | 8 | −3.161 × 10<sup>−6</sup> | 0.003359 | 83.4% | 1167 | 203 |
| 5 h 30 | 7 | 7 | +3.463 × 10<sup>−6</sup> | 0.003278 | 84.2% | 1061 | 182 |
| 24 h | 8 | 5 | +3.161 × 10<sup>−6</sup> | 0.003447 | 82.6% | 977 | 81 |

**Table 2.** Pairwise overlap between the six gene sets. Upper triangle, number of genes in common; lower triangle, the ratio of that number to the chance expectation *n_a_n_b_/N* with *N* = 14 685. Diagonal (italic), the set size *n_a_*. The pair in bold shares the same selected component (*ℓ*_1_ = 5) and is counted as one observation, not two.

|  | 0 min | 45 min | 1 h 30 | 3 h | 5 h 30 | 24 h |
|---|---|---|---|---|---|---|
| 0 min | *1223* | 494 | 565 | 539 | 502 | 485 |
| 45 min | 5.94 | *999* | 500 | 491 | 439 | **633** |
| 1 h 30 | 5.68 | 6.16 | *1194* | 553 | 473 | 490 |
| 3 h | 5.55 | 6.18 | 5.83 | *1167* | 507 | 465 |
| 5 h 30 | 5.68 | 6.08 | 5.48 | 6.01 | *1061* | 433 |
| 24 h | 5.96 | **9.52** | 6.17 | 5.99 | 6.13 | *977* |

**Table 3.** Replication in the 20-day cohort (CRS4). Overlap is counted against the CRS3 gene set at the same time point, over the 14 685 common genes; “chance” is max(*n*_CRS3_*, n*_CRS4_)*/N*, the expected value of (overlap / smaller set).

|  | CRS3 (10 d) |  | CRS4 (20 d) |  | Overlap |  |  |
|---|---|---|---|---|---|---|---|
| Time point | *C* | *n* | *C* | *n* | genes | / smaller | vs. chance |
| 0 min | 87.1% | 1223 | **49.1**% | 371 | 31 | 8.4% | 1.01× |
| 45 min | 84.4% | 999 | 87.4% | 1337 | 504 | 50.5% | 5.55× |
| 1 h 30 | 86.3% | 1194 | 82.3% | 1052 | 460 | 43.7% | 5.38× |
| 3 h | 83.4% | 1167 | 88.8% | 1491 | 597 | 51.2% | 5.04× |
| 5 h 30 | 84.2% | 1061 | 86.8% | 1172 | 476 | 44.9% | 5.62× |
| 24 h | 82.6% | 977 | 86.1% | 1286 | 515 | 52.7% | 6.02× |

Taking 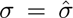 in Eq. (2), *P* -values were corrected by the Benjamini–Hochberg procedure [10] and genes with adjusted *P <* 0.01 were selected. The concentration *C* = 1 *− Nσ*^2^ of Eq. (6) uses the same fitted *σ* and the number of genes entering the tensor (*N* = 14 685 for the RNA-only runs, 13 619 or 13 055 for the six-mode runs). Since 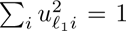 and a background of *N* genes of width *σ* would contribute *Nσ*^2^ to that sum, *C*is the fraction of the squared length of *u_ℓ i_* not attributable to the background; *C* = 0 for an isotropic random unit vector. Counting all *N* genes as background rather than *N* minus the number selected makes *C* a slight underestimate, by 0.8–1.4 percentage points in the runs reported here. *C* does not depend on the significance threshold.

### 2.5 Peak-to-gene aggregation and the six-mode tensor

ATAC peaks (supplied as chr:start-end in Ensembl style) were assigned to genes using windows built from protein-coding transcripts of GENCODE vM17 [11], the annotation used for the RNA quantification, with data.table::foverlaps [12]. Two window definitions were compared: promoter (TSS *±*2 kb) and gene body plus 2 kb upstream, the latter following the GeneActivity definition of Signac [13]. A peak overlapping several windows was counted towards every gene rather than assigned to a nearest one, and chromosome names were stripped of the chr prefix on both sides. The gene body definition was adopted (Table 5). Peaks had been called with Genrich [14] in ATAC mode at FDR 0.05 by the original authors, with mitochondrial reads and ENCODE blacklist regions removed.

**Table 4.** Direction split. The sign of *G* differs between time points, so the mapping from sign(*u*) to direction is not fixed. The two time points in italics select the same component (*ℓ*_1_ = 5) and are strongly unbalanced between the two arms; they are excluded from the comparison below, for the reason given after the table.

| Time point | *n* | same | opposite | same (%) |
|---|---|---|---|---|
| 0 min | 1223 | 602 | 621 | 49 |
| 1 h 30 | 1194 | 649 | 545 | 54 |
| 3 h | 1167 | 684 | 483 | 59 |
| 5 h 30 | 1061 | 648 | 413 | 61 |
| 45 *min* | 999 | 205 | 794 | 21 |
| 24 *h* | 977 | 190 | 787 | 19 |

The six-mode tensor adds an omics index *m ∈ {*1, 2*}* to the brain side only, *z_ijkmsd_* = *x_ijkm_ y_isd_* (Eq. 7), decomposed at ranks (20, 2, 8, 2, 2, 3). RNA, ATAC and skin matrices were each column-normalised independently. The core is then *G*(*ℓ*_1_*ℓ*_2_*ℓ*_3_*ℓ*_4_*ℓ*_5_*ℓ*_6_), and the sign pattern of *u_ℓ_*_4_ *_m_*, the singular vector of the omics mode, labels a component in-phase (uniform sign) or anti-phase (mixed sign).

### 2.6 Enrichment analysis

Gene lists were submitted to the Enrichr REST API [15–17]. The set of 25 libraries was fixed before any result was inspected: GO Biological Process 2026, GO Molecular Function 2026, GO Cellular Component 2026, KEGG 2019 Mouse, Reactome Pathways 2024, MSigDB Hallmark 2020, WikiPathways 2024 Mouse, TRRUST 2019, ChEA 2022, Mouse Gene Atlas, ARCHS4 Tissues, Allen Brain Atlas up, Allen Brain Atlas down, GTEx Tissue Expression Up, GTEx Tissue Expression Down, Tabula Muris, MGI Mammalian Phenotype Level 4 2024, Human Phenotype Ontology, Disease Perturbations from GEO up, Disease Perturbations from GEO down, Jensen DISEASES Curated 2025, DisGeNET, GWAS Catalog 2025, Drug Perturbations from GEO up and Drug Perturbations from GEO down. Libraries assembled automatically from submitted gene lists (Rummagene, RummaGEO, HDSigDB, Enrichr Submissions, Enrichr Users Contributed, PerturbAtlas) were excluded, because they rank highly for any input. Libraries with up and down variants were always included as pairs. Adjusted *P* values and overlap sizes are compared only within a library, never between libraries, since term counts, set sizes and the number of mappable mouse-to-human symbols differ.

### 2.7 Software

All analyses were performed in R with rTensor [9] and data.table [12].

## 3 Results

### 3.1 Construction and selection

The hippocampal dataset [5] profiles the mouse ventral hippocampus by bulk RNA-seq at 0, 45 min, 1 h 30, 3 h, 5 h 30 and 24 h after the initiation of a 1 h 30 restraint, in animals that had previously received 10 or 20 days of daily restraint (chronic restraint stress, CRS) or handling only. Times are measured from the onset of restraint, so that 0 min denotes a home-cage animal with no acute restraint, 45 min an animal still in the tube, and 1 h 30 the moment of release. We used the 10-day cohort (designated CRS3 in the original deposition), contrasting Handling 0min (*n* = 8) with 10daysChronicRestraint at each time point (*n* = 8; *n* = 7 at 5 h 30). Both arms lie within one cohort, so no batch term is needed.

The cutaneous dataset [4] profiles peri-lesional dorsal skin two days after MRSA inoculation, in sham-operated animals that received restraint stress (3 h/day for 3 days) or not (*n* = 3 per group). Adrenalectomised animals from the same series were deliberately left unused, to be available as an independent causal control.

Counts on both sides were transformed as log_2_(*x* + 1) and each sample (column) was scaled to unit norm. Genes were *not* centred and not scaled individually: centring destroys the non-negativity of Eq. (1), after which a product of two negative deviations cannot be distinguished from a product of two positive ones, and in practice no component satisfying the sign conditions then exists (Sec. 3.8). Of 14 689 genes common to both platforms, four Y-linked genes were removed (Sec. 2), leaving *N* = 14 685.

The resulting tensor has shape *N ×* 2 *× K ×* 2 *× D* with *K* = 8 (*K* = 7 at 5 h 30) and *D* = 3, and was decomposed by HOSVD as

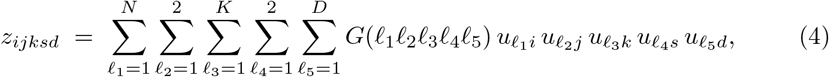

where *G*(*ℓ*_1_*ℓ*_2_*ℓ*_3_*ℓ*_4_*ℓ*_5_) *∈* R*^N×^*^2*×K×*2*×D*^ is the core tensor and the singular-vector matrices are (*u_ℓ i_*) *∈* R*^N×N^*, (*u_ℓ j_*) *∈*R^2*×*2^, (*u_ℓ k_*) *∈* R*^K×K^*, (*u_ℓ s_*) *∈* R^2*×*2^ and (*u_ℓ d_*) *∈*R*^D×D^*, each orthogonal:

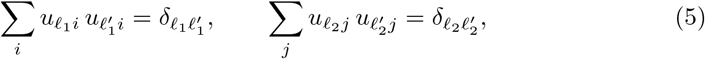

and likewise for modes 3, 4 and 5. In practice the gene mode is truncated at *ℓ*_1_ *≤* 20, so that (*u_ℓ_*_1_ *_i_*) is the 20 *× N* matrix of the leading rows, which is row-orthonormal but not square; modes 2 to 5 are kept at full rank, which makes the truncated decompositions nested and leaves *u_ℓ_*_1_ *_i_* and *G*(*ℓ*_1_*ℓ*_2_*ℓ*_3_*ℓ*_4_*ℓ*_5_) for *ℓ*_1_ *≤* 20 independent of where the gene mode is cut. A component was admitted only if the condition-mode vectors *u_ℓ_*_2_ *_j_* and *u_ℓ_*_4_ *_s_* reverse sign between their two levels and the replicate-mode vectors *u_ℓ_*_3_ *_k_* and *u_ℓ_*_5_ *_d_* are of uniform sign; because the tensor is non-negative, the Perron–Frobenius theorem guarantees that a uniform-sign first singular vector exists in every mode, and the second singular vector of a two-level mode necessarily has mixed signs, so the sign conditions are always satisfiable. Among the admissible components the one with the largest *|G|* along *ℓ*_1_ was taken, and its gene-mode singular vector *u_ℓ_*_1_ *_i_* was fitted to a Gaussian whose width *σ* was chosen to flatten the *P* -value histogram over the null region, and *P* -values were corrected by the Benjamini–Hochberg procedure [10] at 0.01.

### 3.2 A program of the same size is recovered at every time point

Table 1 summarises the six runs. To state how far the gene-mode vector departs from noise we use the concentration

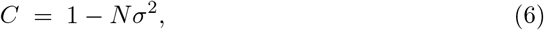

the fraction of the squared length of *u_ℓ_*_1_ *_i_* that is not accounted for by the fitted Gaussian background of width 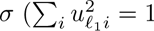, and the *N* genes of a pure background would contribute *σ*^2^ each). *C* = 0 means that *u_ℓ_*_1_ *_i_* is an isotropic random unit vector and nothing has been found; *C* approaching 1 means that almost the whole length of *u_ℓ_*_1_ *_i_* sits in a small number of genes. Equation (6) counts all *N* genes as background and therefore understates *C* slightly; it does not depend on the significance threshold. All six time points fall in the narrow band *C* = 83–87% and return 977–1223 genes.

There is thus no evidence that any one sampling time is privileged. An earlier reading of our own, in which the 3 h result was explained by a match between the hippocampal and cutaneous time constants, is not supported: the structure is present equally at 0 min, when the animal has received no acute restraint at all, and at 24 h, when the original authors detect no differentially expressed genes whatsoever [5].

The *σ* optimisation for a representative time point is shown in Fig. 2; the remaining five are given in the Supplement.

### 3.3 Time points overlap at a common level

The six gene sets overlap one another far more than chance. For sets of size *n_a_* and *n_b_* drawn from *N* = 14 685 genes, the expected overlap if gene identity carried no information is *n_a_n_b_/N*. Table 2 gives all fifteen pairs. Fourteen of them share 433–565 genes, exceeding the expectation by a factor of 5.48–6.18 (Jaccard index 0.265–0.306); 90 genes are common to all six. The spread of the ratio is narrow enough that no pair of time points stands out as more closely related than any other. The single exception, 45 min versus 24 h at 633 genes and 9.52-fold, is not an independent observation, because those two time points select the same component (*ℓ*_1_ = 5).

We stress what this baseline does and does not mean. *n_i_n_j_/N* is the null in which genes are exchangeable. Exceeding it reflects the fact that the same genes are consistently high across samples — the same property that makes cell types identifiable from their highly expressed genes — and would be destroyed by shuffling the gene axis independently within each sample. It is a statement about the data, and we do not read more into it than that.

### 3.4 Replication in an independent cohort

The same deposition contains a second cohort (CRS4) in which animals received 20 rather than 10 days of daily restraint, with its own handling controls. No animal is shared with CRS3, on either arm. We repeated the entire procedure on CRS4 against the same, fixed skin data.

Three readings follow (Table 3).

First, the five acute time points replicate. Every animal is different and the chronic history is twice as long, yet the same genes are selected, at 5.0–6.0 times chance. The identity of the genes is also preserved: at 5 h 30 the CRS4 selection is headed by *Col3a1*, *Col1a1*, *Col1a2*, *Cilp*, *Aebp1*, *Fgfbp1*, *Aqp3*, *Gjb2*, *Cdh1*, *Lox* and *Pcolce* — extracellular matrix and epidermis — together with *Cd74*, *H2-Aa*, *Mrc1*, *Siglec1* and *F13a1*, the same composition as CRS3.

Second, matching the time point buys nothing. The band 5.0–6.0 is the same band as the within-CRS3 overlap between *different* time points (5.48–6.18). Changing the cohort costs as much as changing the time point, and no more. What the procedure recovers is therefore not a time-point-specific response but a core program shared by brain and skin under restraint, reachable from any of the sampling times.

Third, 0 min does not replicate. At 1.01 times chance it is exactly at the null; *C* = 49.1% is the only value outside the 82–89% band spanned by the other eleven runs, and the 371 genes selected — *Thrsp*, *Fosb*, *Krt2*, *Gpd1*, *Mt1*, *Pdk4*, *Dbp*, *Aqp4*, *Cx3cr1*, *Camk2b*, *Snap91*, *Map1a*, *Atp1a3*, *Atp1b2*, *Kcnq2*, *Egr1*, *Nr4a1*, *Ptgs2* — are brain-intrinsic rather than cutaneous. This is a different object, not a weak version of the same one. Whether brain–skin coupling genuinely dissolves at baseline after 20 days of restraint cannot be decided here, because the control animals also differ; we do not pursue it. All claims below are therefore made on the five acute time points, and results that rest on 0 min are flagged as exploratory.

### 3.5 The selected genes belong to both tissues

Enrichment analysis was run through Enrichr [15–17] against a list of 25 libraries fixed in advance of seeing any result (Sec. 2). Across the six time points, 2535 terms are significant at every one of them. Throughout we quote a term as (library, term, range of adjusted *P* across time points, range of overlap size *k*), and we never compare adjusted *P* or *k* between libraries, since term counts, set sizes and the number of mappable genes differ.

The terms discussed below were not chosen after the fact. Table 6 lists, for each of the 25 libraries, the ten most significant terms, ranked by the worst adjusted *P* across the six time points, with no omissions; the complete output is in Additional file 1. Everything we say about enrichment can be checked against that table, including the terms we do not interpret.

Skin is the single strongest attribution in the whole analysis: (ARCHS4 Tissues [18], SKIN (BULK TISSUE), 1.5 *×* 10*^−^*^84^–2.4 *×* 10*^−^*^58^, *k* = 314–379); (ARCHS4 Tissues, KERATINOCYTE, 7.2 *×* 10*^−^*^42^–5.4 *×* 10*^−^*^26^, *k* = 244–303); (Mouse Gene Atlas, epidermis, 1.0 *×* 10*^−^*^36^–2.5 *×* 10*^−^*^24^, *k* = 76–100), the latter being the top term of its library by a margin of fourteen orders of magnitude.

It is not, however, an exclusively cutaneous signature, and Table 6 makes that plain. The ARCHS4 top ten also contains oesophagus, lung, omentum, gastric tissue, colon, breast and ileum, all between 10*^−^*^31^ and 10*^−^*^22^, and the GTEx Tissue Expression Up top ten is occupied not by skin but by oesophagus (1.4 *×* 10*^−^*^42^–1.8 *×* 10*^−^*^25^ for the strongest sample). Likewise, Mouse Gene Atlas places mammary gland, umbilical cord, cornea and osteoblast immediately below epidermis. We read this as it reads: the program is a squamous-epithelial, extracellular-matrix and immune program, of which skin is the strongest but not the only instance. Oesophagus is stratified squamous epithelium like epidermis, and omentum, lung and colon are stroma-rich; a gene set built from keratins, collagens and inflammatory mediators necessarily matches them too. Nothing here argues that the selected genes are specific to skin, and we do not claim that they are.

Attribution to brain is weaker but systematic: (Allen Brain Atlas up [19], 220 regions significant at all six time points); (Allen Brain Atlas down, 16 regions; strongest, layer 6 of occipital cortex, 1.6 *×* 10*^−^*^6^–3.6 *×* 10*^−^*^3^, *k* = 36–47); (GTEx Tissue Expression Down [20], brain samples occupying the whole top ten, 1.0 *×* 10*^−^*^57^–6.2 *×* 10*^−^*^39^, *k* = 260–313). The GTEx “down” direction denotes genes expressed at lower levels in human brain than in other tissues — a statement about tissue specificity in a reference panel, not about the direction of change in our data; the Allen terms likewise denote overlap with genes highly expressed in a region.

Two things about the Allen result should be stated rather than left for a reader to find in Table 6. The brain data are ventral hippocampus, but the regions at the top of Allen Brain Atlas up are not hippocampal: they are brainstem and hypothalamic (lateral lemniscus, medial mammillary nucleus, retromammillary area). And with 220 of the atlas’s regions significant at all six time points, the library has little discriminating power. We therefore use Allen only as evidence that the gene set has measurable overlap with brain-enriched expression at all, and draw no regional conclusion from it.

With that qualification, the gene set is not simply a skin signature that happens to survive the product: it has measurable brain attribution as well, which is what Eq. (1) is constructed to select.

### 3.6 Function: matrix, epithelium and inflammation

The top term of MSigDB Hallmark 2020 is Myogenesis (2.7 *×* 10*^−^*^15^–1.4 *×* 10*^−^*^6^, *k* = 33–47), ahead of every other Hallmark set. We take this to be contamination of mouse dorsal skin by the panniculus carnosus, the subcutaneous striated muscle layer, and we return to it in Sec. 4.3; the driving genes are *Tnnt3*, *Myh4*, *Atp2a1*, *Xirp1/2*, *Ankrd2*, *Myl1* and *Mybpc2*, and the same artefact is visible as cardiac and skeletal muscle entries in Tabula Muris and Mouse Gene Atlas. Setting it aside, the 2535-term core gives a consistent functional picture: (MSigDB Hallmark 2020, Epithelial Mesenchymal Transition, 4.4 *×* 10*^−^*^23^–1.1 *×* 10*^−^*^5^, *k* = 31–53); (Reactome Pathways 2024, Extracellular Matrix Organization, 3.6 *×* 10*^−^*^18^–1.6 *×* 10*^−^*^4^, *k* = 42–61); (MSigDB Hallmark 2020, Interferon Gamma Response, 1.3 *×* 10*^−^*^11^–2.0 *×* 10*^−^*^6^, *k* = 31–40); (MSigDB Hallmark 2020, TNF-alpha Signaling via NF-*κ*B, 6.5 *×* 10*^−^*^14^–4.3 *×* 10*^−^*^4^, *k* = 23–46); (TRRUST 2019 [21], NFKB1 human, 1.1 *×* 10*^−^*^10^–4.9 *×* 10*^−^*^7^, *k* = 43–52); (TRRUST 2019, RELA human, 1.1 *×* 10*^−^*^10^–1.2 *×* 10*^−^*^6^, *k* = 43–54); (GO BP 2026, Inflammatory Response, 7.9 *×* 10*^−^*^12^–1.3 *×* 10*^−^*^5^, *k* = 42–54); (GO BP 2026, Skin Development, 1.9 *×* 10*^−^*^8^–2.5 *×* 10*^−^*^3^, *k* = 14–21); (GO CC 2026, Cornified Envelope, 5.0 *×* 10*^−^*^11^–6.3 *×* 10*^−^*^3^, *k* = 10–18).

At the disease level the picture is mixed, and we state it as the table gives it. In the libraries that are organised by disease category, cutaneous entries lead: (Jensen DISEASES Curated 2025, INTEGUMENTARY SYSTEM DISEASE, 1.9 *×* 10*^−^*^10^–7.2*×*10*^−^*^4^, *k* = 31–48) and (SKIN DISEASE, 2.8*×*10*^−^*^10^–2.1*×*10*^−^*^3^, *k* = 25–42) are that library’s first two terms, followed by KERATOSIS and PALMOPLAN-TAR KERATOSIS, and the Human Phenotype Ontology top ten is almost entirely cutaneous (palmoplantar keratoderma, skin ulcer, plantar hyperkeratosis, erythema, hypohidrosis, blistering). In the libraries built from perturbation experiments, cutaneous entries are present but not dominant. (Disease Perturbations from GEO down, psoriasis DOID-8893, worst 7.3 *×* 10*^−^*^22^) is the top term of that library, and (atopic dermatitis DOID-3310, worst 6.6 *×* 10*^−^*^6^) is also significant throughout; but the corresponding “up” library is headed by acute myocarditis, lung transplant rejection, colitis, pulmonary tuberculosis and cystic fibrosis, with psoriasis (worst 3.7 *×* 10*^−^*^18^) ranked just below them. DisGeNET [22] lists Psoriasis fifth (worst 2.2 *×* 10*^−^*^8^) behind experimental liver cirrhosis, squamous cell carcinoma, asthma and neoplasm metastasis. The common element across these is inflammation with matrix remodelling, in whichever organ the source experiment happened to use; the method cannot distinguish a skin-specific disease signature from a generic one, and we do not claim that it can.

Three libraries in Table 6 carry almost no six-time-point signal at all and should be discounted accordingly. In GO Molecular Function 2026, WikiPathways 2024 Mouse and Human Phenotype Ontology, nine or ten of the top ten terms have a worst adjusted *P* of 1, that is, they fail at one or more time points. GWAS Catalog 2025 has exactly one term significant at all six (Blood Protein Levels, 1.9 *×* 10*^−^*^8^–4.8 *×* 10*^−^*^4^): **there is no evidence here from genetic association**.

The mouse-phenotype library deserves a sentence of its own, because its top ten reads as a description of the cutaneous experiment that generated one half of the tensor: (MGI Mammalian Phenotype Level 4 2024, Abnormal Macrophage Physiology, 2.8 *×* 10*^−^*^12^–1.0 *×* 10*^−^*^3^), (Increased Susceptibility To Induced Morbidity/Mortality, 1.2*×*10*^−^*^7^–1.4*×*10*^−^*^3^), (Abnormal Epidermal Layer Morphology, 4.5*×*10*^−^*^6^–2.1*×*10*^−^*^3^), (Delayed Wound Healing, 2.2 *×* 10*^−^*^7^–2.3 *×* 10*^−^*^3^), (Impaired Skin Barrier Function, 1.2*×*10*^−^*^8^–6.4*×*10*^−^*^3^) and (Increased Susceptibility To Bacterial Infection, 2.2*×*10*^−^*^7^– 9.3 *×* 10*^−^*^3^). Knocking out the genes behind these terms produces, in mice, the phenotype that psychological stress produces in the experiment of Chan et al. [4].

Two coherences in Table 6 run across libraries and were not sought. First, adipogenesis: (TRRUST 2019, PPARG human, 1.7 *×* 10*^−^*^4^–1.6 *×* 10*^−^*^2^) and (PPARG mouse, 4.6 *×* 10*^−^*^4^–2.8 *×* 10*^−^*^2^) are both significant throughout; the top term of Drug Perturbations from GEO down is the PPAR*γ* agonist rosiglitazone (1.4*×*10*^−^*^23^– 9.3 *×* 10*^−^*^14^); and Tabula Muris contributes Mesenchymal Stem Cell Of Adipose Fat (3.6*×*10*^−^*^25^–7.2*×*10*^−^*^8^) alongside Granulocyte Fat. Mouse Gene Atlas adds osteoblast at day 14 and day 21, the other fate of the same mesenchymal progenitors, and RUNX2 — the osteoblast master regulator — is one of the discriminating terms of the opposite-direction arm (Sec. 3.7). Second, TNF: the same drug library’s top ten contains the TNF-blocking biologic etanercept twice, from two independent experiments (1.6 *×* 10*^−^*^17^–1.8 *×* 10*^−^*^9^ and 4.2 *×* 10*^−^*^17^–1.9 *×* 10*^−^*^9^), which is consistent with (Hallmark, TNF-alpha Signaling via NF-*κ*B) and with the NFKB1 and RELA entries above. The fibrosis-inducing agent bleomycin appears three times in the “up” direction of the same library, consistent with the matrix terms.

**Fig. 4.**
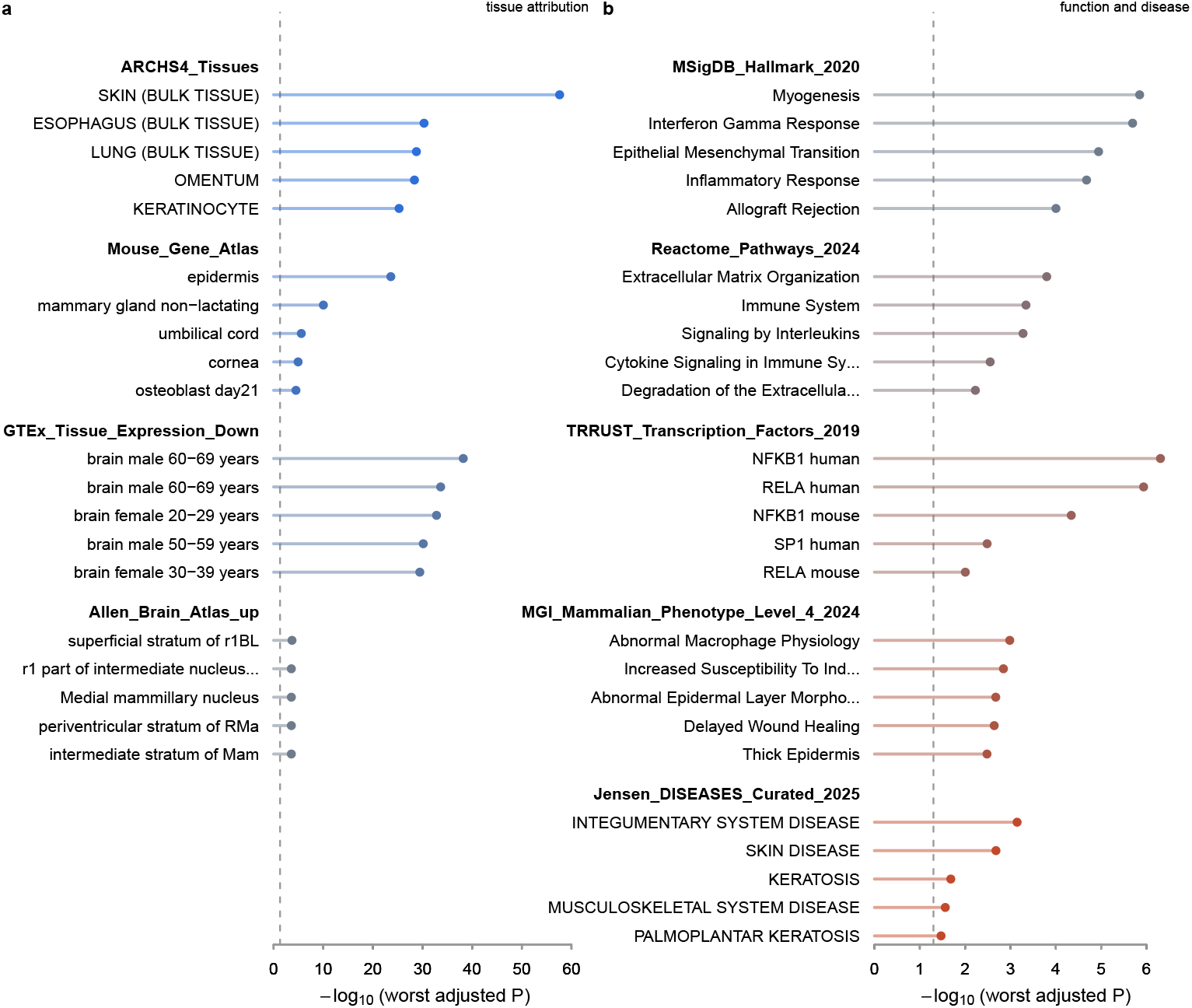
Enrichment across the fixed library set. The five most significant terms of each library, ranked by the worst adjusted *P* across the six time points. (**a**) Tissue attribution. Skin leads ARCHS4 and epidermis leads the Mouse Gene Atlas, but oesophagus, lung and omentum follow closely: the program is squamous-epithelial and stromal rather than exclusively cutaneous. All five leading GTEx samples are brain, in the “down” direction, which denotes genes expressed at lower levels in human brain than in other tissues. (**b**) Function and disease. The dashed line is adjusted *P* = 0.05. Myogenesis heads the Hallmark sets and is most likely contamination by the panniculus carnosus (Sec. 4.3). The full top ten of all 25 libraries is given in Table 6.

Neurological disease terms, by contrast, are *depleted* relative to the library-wide significance rate: 0 observed against 6.6 expected in Jensen DISEASES Curated, 1 against 38.0 in MGI Mammalian Phenotype, and ratios of 0.44–1.01 in DisGeNET and in the two Disease Perturbations libraries, while skin categories run at 1.0–4.7 times the same baseline. We therefore make no claim of brain–skin symmetry at the level of disease. The brain-side evidence rests on expression atlases alone.

Part of this asymmetry is a property of the method. Equation (1) can only select genes that carry a contrast in *both* tissues, and the most neuron-specific genes (*Snap25*, *Syt1*, *Rbfox3*, *Gria1*, *Nefl*, *Slc17a7*, *Gad1*, *Gad2*) are absent from the skin platform and are removed before the tensor is built. Which hippocampal cell types drive the selected genes is therefore not determined by this analysis, and is not a premise of the claim we make.

### 3.7 Same-direction and opposite-direction arms

Because the tensor is non-negative, the sign of *G* together with the sign of *u_ℓ_*_1_ *_i_* fixes the relative direction of the brain and skin contrasts for gene *i*: sign(*G u_ℓ_*_1_ *_i_*) *>* 0 means that the gene moves the same way in both tissues. Table 4 gives the split.

At 45 min and 24 h only 21% and 19% of the selected genes lie on the same-direction side, against 49–61% at the other four. The consequence is visible in the enrichment itself: at those two time points the opposite-direction arm returns 5579 and 5438 significant terms against 800 and 479 on the same-direction side, ratios of 7.0 and 11.4, whereas at the four retained time points the two arms lie within a factor of 1.2 of each other (1.09, 1.10, 1.19 and 1.09). Requiring a term to be significant at every time point of one arm and at none of the other would therefore empty the same-direction class through set size rather than through biology. The two time points were set aside before the comparison was carried out, not after seeing its result; their enrichment was nonetheless run in full and is in Additional file 1 (S2), flagged as not entering the comparison.

Restricting to the four balanced time points, 74 terms are significant on the same-direction side only, 87 on the opposite side only, and 791 on both. Since the same-direction set is the larger one at three of the four time points, any size bias runs against the opposite side, not for it.

The same-direction arm is immune and inflammatory: (Tabula Muris [23], Leukocyte, 1.5 *×* 10*^−^*^7^) is the strongest discriminating term of all; (GTEx Tissue Expression Up, blood samples occupying 47 terms, 2.6 *×* 10*^−^*^5^–8.3 *×* 10*^−^*^4^); (ARCHS4 Tissues, GRANULOCYTE, 2.9 *×* 10*^−^*^2^); (DisGeNET, Inflammatory dermatosis, 4.0 *×* 10*^−^*^4^; Dermatitis, 4.0 *×* 10*^−^*^4^); (GO BP, Positive Regulation of Interleukin-6 Production, 3.3 *×* 10*^−^*^2^); together with epidermal and hair phenotypes (MGI: Abnormal Epidermal Layer Morphology, 1.4 *×* 10*^−^*^2^; Alopecia, 2.9 *×* 10*^−^*^2^). Exactly one brain region appears on this side (Allen Brain Atlas up, medial superior olive, 3.8 *×* 10*^−^*^2^). Adjusted *P* values are the worst of the four time points.

The opposite-direction arm is mesenchymal and vascular: (ChEA 2022, RUNX2, 2.8 *×* 10*^−^*^3^) and (TRRUST, RUNX2 mouse, 4.4 *×* 10*^−^*^2^); (ARCHS4 Tissues, SKELE-TAL MUSCLE SATELLITE CELL, 7.0 *×* 10*^−^*^3^; HEPATIC STELLATE CELL, 2.5 *×* 10*^−^*^2^); (Tabula Muris, Stromal Cell, 3.3 *×* 10*^−^*^3^); (GTEx Up, blood vessel, 2.3 *×* 10*^−^*^3^; adipose tissue, 2.3 *×* 10*^−^*^2^); (GTEx Down, brain, 5.6 *×* 10*^−^*^4^); and eleven Allen brain regions, against one on the same-direction side.

This asymmetry — leukocyte and inflammatory genes moving in concert, matrix and mesenchymal genes moving in opposition — is the single most interpretable structure in the result, and it is reproduced at all four balanced time points independently. Figure 5 summarises the split.

**Fig. 5.**
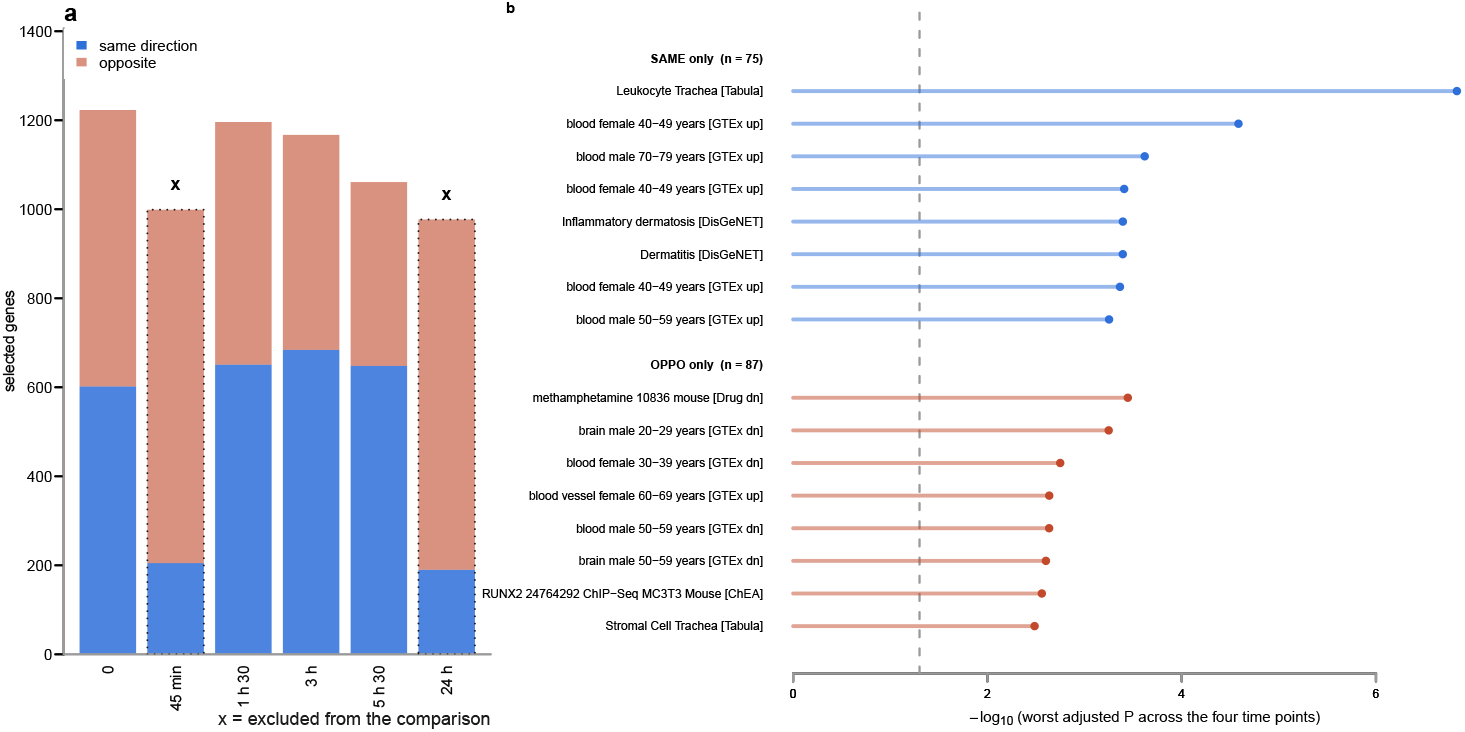
Concordant and discordant arms. (**a**) Selected genes at each time point, split by the sign of *G u_ℓ_*_1_ *_i_*. The two time points marked x are strongly unbalanced between the arms (21% and 19% same-direction) and were excluded from the comparison in advance; their enrichment was run in full and is in Additional file 1. (**b**) Terms significant in one arm only across all four retained time points, the eight strongest of each. The same-direction arm is leukocyte and inflammatory; the opposite-direction arm is mesenchymal and vascular. Library names are abbreviated in brackets.

### 3.8 Controls

#### The procedure returns nothing when there is nothing

Centring the data before forming Eq. (1) destroys non-negativity; no component then satisfies the sign conditions, *C* = 0, and zero genes are selected. The method does not return a thousand genes for any input.

#### Blood contamination is not the explanation

Both tissues contain blood, and stress changes circulating leukocyte composition systemically, so a shared blood signal is a real possibility. If variation in blood content were driving the axis, haematopoietic modules should move together. They do not: at 3 h, *Pf4* (rank 639) and *Hba-a1* (1034) fall inside the selected set while their module partners *Ppbp* (2935), *Itga2b* (3311), *Hbb-bs* (3272), *Alas2* (3127) and *Hba-a2* (7684) fall well outside it. The same argument applies to microglia: *Tyrobp* (578), *P2ry12* (860) and *Aif1* (1094) are selected while *Cx3cr1* (5847), *Tmem119* (6327), *Csf1r* (8069) and *Itgam* (1859) are not. No coherent cell-of-origin story survives this test in either direction.

#### Marginal contrasts do not predict selection

The rank of a gene in *|u_ℓ_*_1_ *_i_|* is not recoverable from its group differences in either dataset. *Sprr2a2*, ranked first at 3 h, has a skin contrast one eighth that of *Grin1*, which ranks 9976th; *Clasrp* (rank 5) and *Arc* (rank 1178) have comparable brain contrasts. The structure exists only in the projected space, which is the reason a joint decomposition is required rather than a comparison of marginal statistics.

#### The gene-mode basis does not depend on the grouping

The mode-1 unfolding of *z* has one column for each pair (((*j, k*), (*s, d*)), equal to the elementwise product of the brain sample (*j, k*) and the skin sample (*s, d*). Reassigning samples between conditions permutes these columns but does not change the column set, so *u_ℓ_*_1_ *_i_* is invariant; what the grouping selects is which of the 20 basis vectors is used. A permutation test on *C* or on the number of selected genes therefore can-not discriminate, and we do not report one. Numerically, 20 composition-preserving regroupings at 3 h gave results that agreed exactly whenever they selected the same component, and the one permutation that selected *ℓ*_1_ = 8 reproduced the observed result exactly.

### 3.9 A second omics layer does not change the result

Bulk ATAC-seq from the same laboratory, the same protocol, the same ventral hippocampi and — critically — the same animals is available for two of the six time points [5]. Rather than analysing it separately, we added it as a fourth mode to the brain side, giving a six-mode tensor

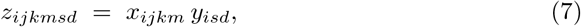

where the brain array now carries an omics index *m ∈ {*1, 2*}* for RNA and ATAC, and *x_ijk_*_1_, *x_ijk_*_2_ are the expression and the aggregated accessibility of gene *i* in the same animal *k* under condition *j*. Because the condition contrast is taken before the omics mode is read, differences in baseline scale between RNA and accessibility cancel, and non-negativity is preserved. Components whose omics-mode singular vector has uniform sign were labelled in-phase and those with opposite signs anti-phase, and both were carried forward in parallel rather than letting max *|G|* choose between them.

Three things follow. Adding accessibility lowers *C* at both time points — from 87.1% to 84.2% at 0 min and from 84.4% to 78.9% at 45 min, with the gene-body windows, and further still with promoter windows — so it does not strengthen the structure. With the gene-body windows, in-phase and anti-phase selected the same component at both time points and therefore returned identical gene sets, so the design did not resolve two populations; with the promoter windows they did differ (813 against 870 genes at 0 min, 791 against 654 at 45 min), but that is the definition we rejected on other grounds. And at 45 min the gene set is essentially unchanged: 497 genes in common, 72.6% of the smaller set against a chance level of 5.9%, with the same leading genes (*Col3a1*, *Emp1*, *Cilp*, *Hp*, *H2-Aa*, *Col1a1*, *Col1a2*, *Cd74*).

**Table 5.** Six-mode tensor with a chromatin-accessibility leg on the brain side. Each cell gives the selected component *ℓ*_1_, the concentration *C* and the number of genes. “RNA only” repeats Table 1 for comparison; note that the common gene set differs because peak-to-gene aggregation does not recover every gene, so *C* is computed with the corresponding *G*.

| Peak assignment | Common genes | 0 min | 45 min |
|---|---|---|---|
| Promoter, TSS ±2 kb | 13 055 | ℓ<sub>1</sub> = 12, 81.4%, 813 | ℓ<sub>1</sub> = 12, 76.5%, 791 |
| Gene body + 2 kb upstream | 13 619 | ℓ<sub>1</sub> = 12, 84.2%, 1139 | ℓ<sub>1</sub> = 8, 78.9%, 685 |
| RNA only | 14 685 | ℓ<sub>1</sub> = 13, 87.1%, 1223 | ℓ<sub>1</sub> = 5, 84.4%, 999 |

This last point is the useful one. The selection is reproduced when an orthogonal measurement of the same animals is admitted into the decomposition, which rules out the possibility that it is an artefact of RNA quantification, library preparation or the particular normalisation applied to counts.

At 0 min the gene set does change substantially (384 genes, 39.6% of the smaller set), and transcription-factor libraries gain disproportionately (TRRUST 22 *→* 94 significant terms; ChEA 80 *→* 179; NFKB1 4.9 *×* 10*^−^*^7^ *→* 9.8 *×* 10*^−^*^11^; RELA 1.2 *×* 10*^−^*^6^ *→* 1.7 *×* 10*^−^*^9^). Since TRRUST is curated from the literature and is not derived from chromatin data, this particular gain cannot be a domain-structure artefact, whereas the ChEA gains for broad-binding factors (SUZ12, SMC3) probably are. We nevertheless record this as exploratory only, because 0 min is the one time point that does not replicate in CRS4 (Sec. 3.4).

## 4 Discussion

### 4.1 What the method contributes

The product tensor of Eq. (1) solves a problem that recurs whenever two tissues are studied by different groups under the same stimulus: the datasets share a feature axis and nothing else. By taking the outer product along that axis, every sample of one study is paired with every sample of the other, so no correspondence has to be invented, no batch term has to be estimated, and the two designs may differ freely in replicate number and platform. Selection then happens in the joint space. A gene that responds strongly in one tissue and not at all in the other receives a small loading, because the product is multiplicative — which is exactly the behaviour one wants from a method intended to find shared programs, and exactly the behaviour that an intersection of two differential-expression lists fails to provide.

Three properties of the construction deserve emphasis. First, non-negativity must be preserved: without it the sign conditions cannot be satisfied and the selection collapses, as the centred control shows. Second, the relative sign of the two contrasts is read directly off sign(*G u_ℓ_*_1_ *_i_*), so the same decomposition that identifies the shared program also partitions it into concordant and discordant arms at no extra cost. Third, the concentration *C* of Eq. (6) provides a calibrated, threshold-free readout of how far the gene-mode vector departs from isotropy, with a known null value of 0 and a maximum of 1.

The method also has a sharp limitation, which our own controls make explicit (Sec. 3.8): the gene-mode basis is invariant under regrouping of samples, so permutation of the labels cannot be used as a null. What the grouping determines is which basis vector is selected, not what the basis is. Validation has to come from independent data, which is why Sec. 3.4 and Sec. 3.9 carry the weight that they do.

### 4.2 What the biology says

The recovered program is simultaneously cutaneous and hippocampal by reference-atlas attribution, and it is dominated by extracellular matrix organisation, epithelial– mesenchymal transition, interferon-*γ* and TNF/NF-*κ*B signalling, with psoriasis as the single strongest disease term in its library.

The direction split is the most informative feature. Leukocyte and inflammatory genes move the same way in brain and skin; matrix, mesenchymal and vascular genes move in opposite directions. A systemic mediator that acts on both tissues — circulating glucocorticoids, or catecholamines — would plausibly produce the concordant arm, and the cutaneous and hippocampal responses to the same signal need not have the same sign in the stromal compartment.

It is worth setting this beside Chan et al. [4], who worked out the cutaneous arm mechanistically: adrenergic signalling induces TGF-*β*, which reprograms dermal fibroblasts undergoing adipogenesis, suppresses cathelicidin and impairs host defence. Three independent features of our result line up with that account, none of which we went looking for. The genes heading the opposite-direction arm are canonical TGF-*β* targets — *Col1a1*, *Col1a2*, *Col3a1*, *Lox*, *Pcolce*, *Aebp1* — and the arm carries stromal, satellite-cell, hepatic-stellate and adipose signatures. The adipogenic programme itself is visible across four unrelated libraries at once: PPAR*γ* in TRRUST, rosiglitazone at the head of the down-regulated drug library, adipose mesenchymal stem cells in Tabula Muris, and osteoblast — the alternative fate of the same progenitor — in Mouse Gene Atlas, with RUNX2 discriminating the opposite-direction arm. And the mouse-phenotype library returns impaired skin barrier function, delayed wound healing and increased susceptibility to bacterial infection, which is their experimental readout stated as a genotype.

We are careful about what this does and does not show. Our analysis was not designed to test their mechanism, contains no perturbation, and cannot establish that TGF-*β* mediates anything. The skin half of the tensor comes from their experiment, so the cutaneous side of these enrichments is not independent evidence about skin. What is new is that the same fibroblast–matrix axis emerges from a decomposition in which the hippocampal contrast is an equal partner and in which no cutaneous contrast alone can select a gene, and that it emerges as the *discordant* arm — moving oppositely in the two tissues — while the leukocyte arm moves concordantly.

Against this, the brain side of the result is weaker than the skin side and we say so plainly. Neurological disease and phenotype terms are depleted, not enriched, and the hippocampal attribution rests on expression atlases alone. Part of this is structural — the most neuron-specific genes never enter the common gene set — but we cannot exclude that the hippocampal contribution is carried largely by non-neuronal and vascular compartments, and nothing in this analysis determines the cell of origin.

Finally, the finding that matching the time point confers no advantage over matching nothing (Sec. 3.4) is itself a result about the biology. The program is not a transient wave that the skin and the brain happen to ride simultaneously; it is a standing relationship between the two tissues under restraint, detectable from any point in the 24 h following the stressor, including 24 h, where the original study finds no differentially expressed genes at all.

### 4.3 Limitations

The cutaneous arm is never varied. The same six skin samples enter all twelve runs, so the replication of Sec. 3.4 establishes independence of the hippocampal animals only, and the overlap figures do not contain cutaneous sampling variance. No second dataset pairing brain and skin under a common stressor exists in public repositories, so this cannot presently be remedied.

The two studies differ in sex composition: the hippocampal experiment used male and female mice in equal numbers, while the cutaneous experiment used males only. Y-linked genes were removed before the tensor was built, and replicate pairs were matched by sex on the brain side, but a sex-dependent component of the cutaneous response cannot be assessed.

The projection of samples onto the selected axis separates the groups, but that axis was derived from the same data and the separation is constructive, not an independent test; we therefore do not report it as evidence. The glucocorticoid reading of the direction split is post hoc and does not fall cleanly on one side: dexamethasone appears in the down-regulated drug library, but so do rosiglitazone, formaldehyde, sunitinib, phorbol ester and sodium arsenite, and the up direction of that library is headed by soman, which is not interpretable here at all.

A myogenic signature is the top Hallmark term (2.7 *×* 10*^−^*^15^–1.4 *×* 10*^−^*^6^; *Tnnt3*, *Myh4*, *Atp2a1*, *Xirp1/2*, *Ankrd2*, *Myl1*, *Mybpc2*), and reappears as cardiac and skeletal muscle entries in Tabula Muris and Mouse Gene Atlas and as impaired skeletal muscle contractility in MGI. The most likely explanation is contamination of mouse dorsal skin by the panniculus carnosus, the subcutaneous striated muscle layer, which would be dissected with the skin. We cannot exclude it, and we do not interpret any muscle term.

The tissue attribution is not specific to skin. Oesophagus, lung, omentum, gastric tissue, colon and breast accompany skin in the ARCHS4 top ten, and oesophagus rather than skin heads GTEx Tissue Expression Up. The program should be read as squamous-epithelial, matrix and immune, not as cutaneous in the exclusive sense.

Finally, *Staphylococcus aureus* infection appears as a KEGG term, and the cutaneous half of the tensor comes from infected animals. The brain–skin program we describe is defined under that infection and may not generalise to uninfected skin; no public dataset lets us check this.

## 5 Conclusions

Two mouse studies that share a stressor and a gene list, and nothing else — not animals, not laboratory, not sequencing run — turn out to share a transcriptional program. Multiplying them into a single five-mode tensor and decomposing it returns between 977 and 1223 genes at each of six time points, with 82.6–87.1% of the squared length of the gene-mode vector lying outside the fitted null. An entirely separate cohort of animals, chronically stressed for twenty days instead of ten, recovers the same genes at 5.04 to 6.02 times chance at five of those six time points — the same band as the agreement between time points within one cohort, so nothing here depends on matching the hour. Adding chromatin accessibility as a sixth mode leaves the gene set essentially unchanged.

The program is squamous-epithelial, matrix and immune, and it separates by direction: leukocyte and inflammatory genes move together in the two tissues, matrix and mesenchymal genes move against each other. That asymmetry, not the gene list, is the result we would most want tested next.

What makes this possible is that the product tensor needs no correspondence between the two datasets. Wherever two tissues have been profiled by different groups under one stimulus — the ordinary case in public data — the same construction applies, and the question of whether they share a response becomes answerable without new experiments. The brain–skin case would still be settled properly by one study sampling both tissues in the same animals; until such a dataset exists, an analysis of this kind is what the public record allows.

## Supplementary information

### Additional files

**Additional file 1.** SupplementaryData1.zip (ZIP archive; six comma-separated-value files, one of them gzip compressed, and a directory of plain-text gene lists). *Complete enrichment output and gene lists.* Additional file 1 contains the complete output of the analysis, so that every enrichment statement in this paper can be checked and the selected genes recovered. Nothing in it has been filtered by hand. It holds six parts:

*S1* Every term of all 25 libraries at the six time points (9396 rows), with adjusted *P* and *k* per time point and the worst and best adjusted *P* across them. Table 6 is the first ten rows of each library after sorting this file.

**Table 6:** The ten most significant terms of every library, ranked by the worst adjusted *P* across the six time points. Within a library’s top ten nothing is omitted. *k*, number of genes in the overlap; the two *P* values and the *k* range are taken over the six time points. Adjusted *P* and *k* are comparable within a library but not between libraries.

| Term | worst adj. *P* | best adj. *P* | *k* |
|---|---|---|---|
| **GO_Biological_Process_2026** |  |  |  |
| Inflammatory Response (GO:0006954) | 1.3 × 10<sup>−5</sup> | 7.9 × 10<sup>−12</sup> | 42–54 |
| Extracellular Matrix Organization (GO:0030198) | 6.0 × 10<sup>−5</sup> | 3.9 × 10<sup>−11</sup> | 30–41 |
| Regulation of Angiogenesis (GO:0045765) | 1.4 × 10<sup>−3</sup> | 2.4 × 10<sup>−7</sup> | 28–39 |
| Skin Development (GO:0043588) | 2.5 × 10<sup>−3</sup> | 1.9 × 10<sup>−8</sup> | 14–21 |
| Regulation of Inflammatory Response (GO:0050727) | 3.2 × 10<sup>−3</sup> | 3.9 × 10<sup>−10</sup> | 31–49 |
| Epidermis Development (GO:0008544) | 3.3 × 10<sup>−3</sup> | 3.2 × 10<sup>−7</sup> | 16–24 |
| Cytokine-mediated Signaling Pathway (GO:0019221) | 4.0 × 10<sup>−3</sup> | 5.5 × 10<sup>−7</sup> | 31–47 |
| Positive Regulation of Cell Migration (GO:0030335) | 8.5 × 10<sup>−3</sup> | 1.4 × 10<sup>−6</sup> | 34–52 |
| Positive Regulation of Angiogenesis (GO:0045766) | 9.3 × 10<sup>−3</sup> | 2.6 × 10<sup>−4</sup> | 17–23 |
| External Encapsulating Structure Organization (GO:0045229) | 1.5 × 10<sup>−2</sup> | 5.0 × 10<sup>−7</sup> | 16–24 |
| **GO_Molecular_Function_2026** |  |  |  |
| Cytokine Receptor Activity (GO:0004896) | 1.5 × 10<sup>−2</sup> | 5.1 × 10<sup>−10</sup> | 15–27 |
| Serine-type Peptidase Activity (GO:0008236) | ≥ 1 | 1.4 × 10<sup>−5</sup> | 0–28 |
| Serine-type Endopeptidase Activity (GO:0004252) | ≥ 1 | 4.0 × 10<sup>−5</sup> | 0–24 |
| Calcium Ion Binding (GO:0005509) | ≥ 1 | 4.5 × 10<sup>−4</sup> | 0–39 |
| Tropomyosin Binding (GO:0005523) | ≥ 1 | 4.5 × 10<sup>−4</sup> | 0–7 |
| Chemokine Receptor Binding (GO:0042379) | ≥ 1 | 1.2 × 10<sup>−3</sup> | 0–14 |
| Exopeptidase Activity (GO:0008238) | ≥ 1 | 1.4 × 10<sup>−3</sup> | 0–12 |
| Endopeptidase Inhibitor Activity (GO:0004866) | ≥ 1 | 1.5 × 10<sup>−3</sup> | 0–19 |
| Chemokine Activity (GO:0008009) | ≥ 1 | 1.8 × 10<sup>−3</sup> | 0–13 |
| C-C Chemokine Binding (GO:0019957) | ≥ 1 | 2.2 × 10<sup>−3</sup> | 0–8 |
| **GO_Cellular_Component_2026** |  |  |  |
| Cornified Envelope (GO:0001533) | 6.3 × 10<sup>−3</sup> | 5.0 × 10<sup>−11</sup> | 10–18 |
| Supramolecular Fiber (GO:0099512) | 2.5 × 10<sup>−2</sup> | 1.9 × 10<sup>−16</sup> | 12–27 |
| Secretory Granule Lumen (GO:0034774) | 2.5 × 10<sup>−2</sup> | 9.5 × 10<sup>−4</sup> | 32–38 |
| Desmosome (GO:0030057) | 3.0 × 10<sup>−2</sup> | 3.8 × 10<sup>−7</sup> | 6–11 |
| Extracellular Exosome (GO:0070062) | 3.3 × 10<sup>−2</sup> | 7.4 × 10<sup>−14</sup> | 140–183 |
| Extracellular Vesicle (GO:1903561) | 3.3 × 10<sup>−2</sup> | 1.1 × 10<sup>−13</sup> | 142–184 |
| Elastic Fiber (GO:0071953) | 4.4 × 10<sup>−2</sup> | 2.6 × 10<sup>−15</sup> | 8–21 |
| Intermediate Filament Cytoskeleton (GO:0045111) | ≥ 1 | 3.0 × 10<sup>−6</sup> | 0–16 |
| Endoplasmic Reticulum Lumen (GO:0005788) | ≥ 1 | 2.1 × 10<sup>−5</sup> | 0–42 |
| Intermediate Filament (GO:0005882) | ≥ 1 | 6.6 × 10<sup>−5</sup> | 0–17 |
| **KEGG_2019_Mouse** |  |  |  |
| Cytokine-cytokine receptor interaction | 1.9 × 10<sup>−3</sup> | 2.2 × 10<sup>−10</sup> | 30–53 |
| Proteoglycans in cancer | 9.2 × 10<sup>−3</sup> | 1.9 × 10<sup>−6</sup> | 21–35 |
| ECM-receptor interaction | 1.2 × 10<sup>−2</sup> | 1.5 × 10<sup>−6</sup> | 13–19 |
| Phagosome | 1.4 × 10<sup>−2</sup> | 4.3 × 10<sup>−14</sup> | 20–41 |
| Rheumatoid arthritis | 1.4 × 10<sup>−2</sup> | 6.2 × 10<sup>−4</sup> | 12–16 |
| Human T-cell leukemia virus 1 infection | 1.5 × 10<sup>−2</sup> | 4.5 × 10<sup>−4</sup> | 27–31 |
| AGE-RAGE signaling pathway in diabetic complications | 1.9 × 10<sup>−2</sup> | 9.0 × 10<sup>−5</sup> | 13–20 |
| Tuberculosis | 1.9 × 10<sup>−2</sup> | 4.5 × 10<sup>−5</sup> | 19–29 |
| Toxoplasmosis | 2.0 × 10<sup>−2</sup> | 3.6 × 10<sup>−4</sup> | 14–20 |
| Malaria | 2.4 × 10<sup>−2</sup> | 1.2 × 10<sup>−6</sup> | 8–15 |
| **Reactome_Pathways_2024** |  |  |  |
| Extracellular Matrix Organization | 1.6 × 10<sup>−4</sup> | 3.6 × 10<sup>−18</sup> | 42–61 |
| Immune System | 4.6 × 10<sup>−4</sup> | 3.8 × 10<sup>−8</sup> | 161–186 |
| Signaling by Interleukins | 5.3 × 10<sup>−4</sup> | 4.8 × 10<sup>−7</sup> | 50–57 |
| Cytokine Signaling in Immune System | 2.8 × 10<sup>−3</sup> | 2.3 × 10<sup>−7</sup> | 73–84 |
| Degradation of the Extracellular Matrix | 5.8 × 10<sup>−3</sup> | 6.4 × 10<sup>−11</sup> | 21–32 |
| Neutrophil Degranulation | 8.4 × 10<sup>−3</sup> | 1.2 × 10<sup>−5</sup> | 45–61 |
| Interleukin-4 and Interleukin-13 Signaling | 1.5 × 10<sup>−2</sup> | 1.1 × 10<sup>−6</sup> | 17–23 |
| Interleukin-10 Signaling | 2.8 × 10<sup>−2</sup> | 1.5 × 10<sup>−4</sup> | 9–14 |
| Assembly of Collagen Fibrils and Other Multimeric Structures | 3.6 × 10<sup>−2</sup> | 1.0 × 10<sup>−8</sup> | 11–19 |
| Chemokine Receptors Bind Chemokines | 3.9 × 10<sup>−2</sup> | 4.0 × 10<sup>−5</sup> | 9–17 |
| **MSigDB_Hallmark_2020** |  |  |  |
| Myogenesis | 1.4 × 10<sup>−6</sup> | 2.7 × 10<sup>−15</sup> | 33–47 |
| Interferon Gamma Response | 2.0 × 10<sup>−6</sup> | 1.3 × 10<sup>−11</sup> | 31–40 |
| Epithelial Mesenchymal Transition | 1.1 × 10<sup>−5</sup> | 4.4 × 10<sup>−23</sup> | 31–53 |
| Inflammatory Response | 2.1 × 10<sup>−5</sup> | 2.2 × 10<sup>−9</sup> | 28–38 |
| Allograft Rejection | 9.9 × 10<sup>−5</sup> | 6.6 × 10<sup>−9</sup> | 25–35 |
| TNF-alpha Signaling via NF-kB | 4.3 × 10<sup>−4</sup> | 6.5 × 10<sup>−14</sup> | 23–46 |
| KRAS Signaling Up | 8.2 × 10<sup>−4</sup> | 1.8 × 10<sup>−10</sup> | 24–40 |
| IL-6/JAK/STAT3 Signaling | 2.4 × 10<sup>−3</sup> | 3.7 × 10<sup>−8</sup> | 13–22 |
| Apoptosis | 3.2 × 10<sup>−3</sup> | 1.5 × 10<sup>−5</sup> | 19–26 |
| Interferon Alpha Response | 4.2 × 10<sup>−3</sup> | 1.1 × 10<sup>−7</sup> | 14–21 |
| **WikiPathways_2024_Mouse** |  |  |  |
| Lung Fibrosis WP3632 | ≥ 1 | 9.8 × 10<sup>−6</sup> | 0–17 |
| Tyrobp Causal Network In Microglia WP3625 | ≥ 1 | 1.3 × 10<sup>−4</sup> | 0–15 |
| Prostaglandin Synthesis And Regulation WP374 | ≥ 1 | 1.6 × 10<sup>−4</sup> | 0–10 |
| Macrophage Markers WP2271 | ≥ 1 | 5.3 × 10<sup>−4</sup> | 0–6 |
| Spinal Cord Injury WP2432 | ≥ 1 | 5.6 × 10<sup>−4</sup> | 0–19 |
| Focal Adhesion WP85 | ≥ 1 | 5.6 × 10<sup>−4</sup> | 0–25 |
| Eicosanoid Lipid Synthesis Map WP4335 | ≥ 1 | 1.2 × 10<sup>−3</sup> | 0–5 |
| Focal Adhesion PI3K Akt mTOR Signaling Pathway WP2841 | ≥ 1 | 1.9 × 10<sup>−3</sup> | 0–38 |
| Endochondral Ossification WP1270 | ≥ 1 | 1.9 × 10<sup>−3</sup> | 0–13 |
| Inflammatory Response Pathway WP458 | ≥ 1 | 2.2 × 10<sup>−3</sup> | 0–8 |
| **TRRUST_Transcription_Factors_2019** |  |  |  |
| NFKB1 human | 4.9 × 10<sup>−7</sup> | 1.1 × 10<sup>−10</sup> | 43–52 |
| RELA human | 1.2 × 10<sup>−6</sup> | 1.1 × 10<sup>−10</sup> | 43–54 |
| NFKB1 mouse | 4.6 × 10<sup>−5</sup> | 8.2 × 10<sup>−9</sup> | 30–37 |
| SP1 human | 3.3 × 10<sup>−3</sup> | 6.9 × 10<sup>−6</sup> | 46–58 |
| RELA mouse | 1.0 × 10<sup>−2</sup> | 2.0 × 10<sup>−3</sup> | 16–19 |
| PPARG human | 1.6 × 10<sup>−2</sup> | 1.7 × 10<sup>−4</sup> | 11–14 |
| TRP53 mouse | 2.6 × 10<sup>−2</sup> | 1.5 × 10<sup>−4</sup> | 19–26 |
| STAT1 human | 2.7 × 10<sup>−2</sup> | 1.6 × 10<sup>−4</sup> | 12–18 |
| PPARG mouse | 2.8 × 10<sup>−2</sup> | 4.6 × 10<sup>−4</sup> | 8–13 |
| SP1 mouse | 4.5 × 10<sup>−2</sup> | 6.9 × 10<sup>−6</sup> | 28–39 |
| **ChEA_2022** |  |  |  |
| P63 26484246 Chip-Seq KERATINOCYTES Human | 1.0 × 10<sup>−5</sup> | 3.0 × 10<sup>−16</sup> | 136–181 |
| SOX2 20726797 ChIP-Seq SW620 Human | 6.2 × 10<sup>−5</sup> | 2.2 × 10<sup>−11</sup> | 151–195 |
| RELA 24523406 ChIP-Seq FIBROSARCOMA Human | 1.4 × 10<sup>−4</sup> | 8.0 × 10<sup>−8</sup> | 95–111 |
| STAT5 23275557 ChIP-Seq MAMMARY-EPITHELIUM Mouse | 2.4 × 10<sup>−4</sup> | 3.3 × 10<sup>−7</sup> | 82–98 |
| PBX 27287812 Chip-Seq EMBYONIC-LIMB Mouse | 2.7 × 10<sup>−4</sup> | 7.5 × 10<sup>−9</sup> | 119–151 |
| BACH1 22875853 ChIP-PCR HELA AND SCP4 Human | 2.8 × 10<sup>−4</sup> | 6.9 × 10<sup>−11</sup> | 96–117 |
| SMAD2 18955504 ChIP-ChIP HaCaT Human | 4.0 × 10<sup>−4</sup> | 5.0 × 10<sup>−8</sup> | 116–142 |
| SMAD3 18955504 ChIP-ChIP HaCaT Human | 4.0 × 10<sup>−4</sup> | 5.0 × 10<sup>−8</sup> | 116–142 |
| TCF21 26020271 ChIP-Seq SMOOTH MUSCLE Human | 5.0 × 10<sup>−4</sup> | 1.1 × 10<sup>−11</sup> | 133–155 |
| IRF8 27001747 Chip-Seq BMDM Mouse | 5.7 × 10<sup>−4</sup> | 1.0 × 10<sup>−9</sup> | 108–159 |
| **Mouse_Gene_Atlas** |  |  |  |
| epidermis | 2.5 × 10<sup>−24</sup> | 1.0 × 10<sup>−36</sup> | 76–100 |
| mammary gland non-lactating | 1.0 × 10<sup>−10</sup> | 7.3 × 10<sup>−18</sup> | 37–51 |
| umbilical cord | 2.7 × 10<sup>−6</sup> | 6.4 × 10<sup>−12</sup> | 31–43 |
| cornea | 1.1 × 10<sup>−5</sup> | 4.4 × 10<sup>−10</sup> | 50–62 |
| osteoblast day21 | 3.2 × 10<sup>−5</sup> | 1.2 × 10<sup>−22</sup> | 35–61 |
| stomach | 5.4 × 10<sup>−4</sup> | 4.5 × 10<sup>−7</sup> | 28–40 |
| osteoblast day14 | 1.1 × 10<sup>−2</sup> | 2.9 × 10<sup>−16</sup> | 29–56 |
| intestine large | 3.5 × 10<sup>−2</sup> | 6.3 × 10<sup>−4</sup> | 30–41 |
| bladder | ≥ 1 | 3.8 × 10<sup>−13</sup> | 0–40 |
| skeletal muscle | ≥ 1 | 3.8 × 10<sup>−6</sup> | 0–71 |
| **ARCHS4_Tissues** |  |  |  |
| SKIN (BULK TISSUE) | 2.4 × 10<sup>−58</sup> | 1.5 × 10<sup>−84</sup> | 314–379 |
| ESOPHAGUS (BULK TISSUE) | 5.0 × 10<sup>−31</sup> | 7.9 × 10<sup>−54</sup> | 257–318 |
| LUNG (BULK TISSUE) | 1.7 × 10<sup>−29</sup> | 2.8 × 10<sup>−63</sup> | 253–341 |
| OMENTUM | 4.3 × 10<sup>−29</sup> | 4.4 × 10<sup>−72</sup> | 259–327 |
| KERATINOCYTE | 5.4 × 10<sup>−26</sup> | 7.2 × 10<sup>−42</sup> | 244–303 |
| GASTRIC TISSUE (BULK) | 2.6 × 10<sup>−25</sup> | 3.2 × 10<sup>−50</sup> | 242–313 |
| COLON (BULK TISSUE) | 5.5 × 10<sup>−25</sup> | 1.0 × 10<sup>−47</sup> | 241–290 |
| BREAST (BULK TISSUE) | 2.7 × 10<sup>−24</sup> | 1.1 × 10<sup>−48</sup> | 239–301 |
| AMNIOTIC FLUID | 1.3 × 10<sup>−23</sup> | 2.0 × 10<sup>−31</sup> | 237–274 |
| ILEUM (BULK) | 1.5 × 10<sup>−22</sup> | 4.2 × 10<sup>−39</sup> | 234–288 |
| **Allen_Brain_Atlas_up** |  |  |  |
| superficial stratum of r1BL | 2.0 × 10<sup>−4</sup> | 1.5 × 10<sup>−12</sup> | 37–52 |
| r1 part of intermediate nucleus of the lateral lemniscus | 2.7 × 10<sup>−4</sup> | 1.5 × 10<sup>−12</sup> | 36–52 |
| Medial mammillary nucleus | 2.7 × 10<sup>−4</sup> | 1.6 × 10<sup>−11</sup> | 43–59 |
| periventricular stratum of RMa | 2.7 × 10<sup>−4</sup> | 1.5 × 10<sup>−10</sup> | 36–49 |
| intermediate stratum of Mam | 2.7 × 10<sup>−4</sup> | 3.8 × 10<sup>−9</sup> | 36–45 |
| retromammillary area, lateral part | 2.7 × 10<sup>−4</sup> | 9.7 × 10<sup>−8</sup> | 35–49 |
| intermediate stratum of RMa | 2.7 × 10<sup>−4</sup> | 1.5 × 10<sup>−7</sup> | 34–48 |
| Nucleus of the lateral lemniscus, ventral part | 4.3 × 10<sup>−4</sup> | 1.4 × 10<sup>−13</sup> | 35–54 |
| retromammillary area, medial part | 4.3 × 10<sup>−4</sup> | 1.5 × 10<sup>−10</sup> | 35–51 |
| Nucleus of the lateral lemniscus, horizontal part | 4.3 × 10<sup>−4</sup> | 1.3 × 10<sup>−9</sup> | 35–46 |
| **Allen_Brain_Atlas_down** |  |  |  |
| layer 6 of OCx | 3.6 × 10<sup>−3</sup> | 1.6 × 10<sup>−6</sup> | 36–47 |
| r2 part of nucleus subcoeruleus | 1.2 × 10<sup>−2</sup> | 1.6 × 10<sup>−6</sup> | 32–42 |
| Primary somatosensory area, lower limb, layer 6a | 1.8 × 10<sup>−2</sup> | 3.5 × 10<sup>−4</sup> | 31–36 |
| liminal part of r3 alar plate | 1.9 × 10<sup>−2</sup> | 1.1 × 10<sup>−3</sup> | 30–36 |
| Primary somatosensory area, trunk, layer 6b | 2.1 × 10<sup>−2</sup> | 4.6 × 10<sup>−5</sup> | 31–43 |
| sublayer 6b of FCx | 2.5 × 10<sup>−2</sup> | 2.7 × 10<sup>−5</sup> | 32–44 |
| lateral wings of the dorsal raphe nucleus | 2.5 × 10<sup>−2</sup> | 3.7 × 10<sup>−4</sup> | 27–40 |
| Anteromedial visual area, layer 6a | 2.9 × 10<sup>−2</sup> | 1.9 × 10<sup>−6</sup> | 30–47 |
| r1 part of the ’mesencephalic’ trigeminal nucleus | 3.0 × 10<sup>−2</sup> | 6.3 × 10<sup>−5</sup> | 27–43 |
| layer 5 of CCx | 3.0 × 10<sup>−2</sup> | 1.5 × 10<sup>−4</sup> | 31–36 |
| **GTEx_Tissue_Expression_Up** |  |  |  |
| GTEX-WFJO-0826-SM-4LVM5 esophagus male 30-39 years | 1.8 × 10<sup>−25</sup> | 1.4 × 10<sup>−42</sup> | 158–214 |
| GTEX-Q2AI-0626-SM-48TZU esophagus male 40-49 years | 6.0 × 10<sup>−25</sup> | 1.3 × 10<sup>−35</sup> | 140–171 |
| GTEX-SUCS-0726-SM-4DM7J esophagus male 70-79 years | 1.2 × 10<sup>−24</sup> | 1.2 × 10<sup>−31</sup> | 140–170 |
| GTEX-RM2N-0726-SM-48FD5 esophagus male 50-59 years | 7.9 × 10<sup>−24</sup> | 2.6 × 10<sup>−35</sup> | 149–176 |
| GTEX-XXEK-0326-SM-4BRVV esophagus male 50-59 years | 2.0 × 10<sup>−23</sup> | 1.5 × 10<sup>−32</sup> | 155–193 |
| GTEX-QLQ7-0926-SM-447BC esophagus male 60-69 years | 3.2 × 10<sup>−23</sup> | 2.8 × 10<sup>−35</sup> | 143–181 |
| GTEX-U3ZN-1626-SM-4DXTZ cervix uteri female 30-39 years | 9.8 × 10<sup>−23</sup> | 1.6 × 10<sup>−35</sup> | 162–205 |
| GTEX-SNMC-0526-SM-4DM69 esophagus male 20-29 years | 2.8 × 10<sup>−22</sup> | 7.7 × 10<sup>−34</sup> | 127–161 |
| GTEX-T2IS-0826-SM-4DM6L esophagus female 20-29 years | 1.0 × 10<sup>−21</sup> | 2.1 × 10<sup>−33</sup> | 149–197 |
| GTEX-VUSG-1226-SM-4KKZF esophagus male 50-59 years | 1.0 × 10<sup>−21</sup> | 3.7 × 10<sup>−31</sup> | 138–169 |
| **GTEx_Tissue_Expression_Down** |  |  |  |
| GTEX-N7MS-0011-R11A-SM-2HMJS brain male 60-69 years | 6.2 × 10<sup>−39</sup> | 1.0 × 10<sup>−57</sup> | 260–313 |
| GTEX-N7MS-2526-SM-2D7W3 brain male 60-69 years | 2.2 × 10<sup>−34</sup> | 2.1 × 10<sup>−48</sup> | 230–270 |
| GTEX-T2IS-0011-R11A-SM-32QPC brain female 20-29 years | 1.5 × 10<sup>−33</sup> | 3.4 × 10<sup>−68</sup> | 229–291 |
| GTEX-WVLH-0011-R11A-SM-3MJFO brain male 50-59 years | 7.1 × 10<sup>−31</sup> | 6.1 × 10<sup>−53</sup> | 222–281 |
| GTEX-QDT8-3026-SM-32PKB brain female 30-39 years | 3.5 × 10<sup>−30</sup> | 3.3 × 10<sup>−41</sup> | 202–237 |
| GTEX-WL46-2826-SM-3LK81 brain male 50-59 years | 4.9 × 10<sup>−30</sup> | 2.2 × 10<sup>−52</sup> | 231–296 |
| GTEX-S7SE-2526-SM-2XCDL brain male 50-59 years | 1.1 × 10<sup>−29</sup> | 1.2 × 10<sup>−48</sup> | 180–227 |
| GTEX-RVPV-0011-R11A-SM-2TF6F brain male 60-69 years | 4.9 × 10<sup>−28</sup> | 4.8 × 10<sup>−42</sup> | 225–271 |
| GTEX-T6MN-0011-R11A-SM-32QOX brain male 50-59 years | 1.9 × 10<sup>−27</sup> | 3.7 × 10<sup>−48</sup> | 197–252 |
| GTEX-QMR6-0011-R11A-SM-32PKK brain male 50-59 years | 8.7 × 10<sup>−25</sup> | 3.7 × 10<sup>−41</sup> | 211–262 |
| **Tabula_Muris** |  |  |  |
| Macrophage Muscle CL:0000235 | 6.1 × 10<sup>−9</sup> | 4.0 × 10<sup>−15</sup> | 20–28 |
| Epidermal Cell Skin CL:0000362 | 6.1 × 10<sup>−9</sup> | 3.5 × 10<sup>−13</sup> | 20–24 |
| Monocyte Marrow CL:0000576 | 1.3 × 10<sup>−8</sup> | 2.4 × 10<sup>−13</sup> | 20–26 |
| Granulocyte Fat CL:0000094 | 2.0 × 10<sup>−8</sup> | 2.5 × 10<sup>−16</sup> | 20–29 |
| Mesenchymal Stem Cell Muscle CL:0000134 | 2.3 × 10<sup>−8</sup> | 2.7 × 10<sup>−18</sup> | 23–32 |
| Fibroblast Heart CL:0000057 | 3.2 × 10<sup>−8</sup> | 8.1 × 10<sup>−25</sup> | 23–39 |
| Cardiac Muscle Cell Heart CL:0000746 | 3.4 × 10<sup>−8</sup> | 3.9 × 10<sup>−18</sup> | 20–35 |
| Leukocyte Lung CL:0000738 | 4.2 × 10<sup>−8</sup> | 2.7 × 10<sup>−17</sup> | 18–30 |
| Leukocyte Kidney CL:0000738 | 5.3 × 10<sup>−8</sup> | 1.9 × 10<sup>−11</sup> | 18–23 |
| Mesenchymal Stem Cell Of Adipose Fat CL:0002570 | 7.2 × 10<sup>−8</sup> | 3.6 × 10<sup>−25</sup> | 22–39 |
| **MGI_Mammalian_Phenotype_Level_4_2024** |  |  |  |
| Abnormal Macrophage Physiology MP:0002451 | 1.0 × 10<sup>−3</sup> | 2.8 × 10<sup>−12</sup> | 26–44 |
| Increased Susceptibility To Induced Morbidity/Mortality MP:0009763 | 1.4 × 10<sup>−3</sup> | 1.2 × 10<sup>−7</sup> | 28–35 |
| Abnormal Epidermal Layer Morphology MP:0001216 | 2.1 × 10<sup>−3</sup> | 4.5 × 10<sup>−6</sup> | 13–19 |
| Delayed Wound Healing MP:0002908 | 2.3 × 10<sup>−3</sup> | 2.2 × 10<sup>−7</sup> | 12–18 |
| Thick Epidermis MP:0001219 | 3.3 × 10<sup>−3</sup> | 3.8 × 10<sup>−5</sup> | 13–18 |
| Decreased Inflammatory Response MP:0001876 | 4.0 × 10<sup>−3</sup> | 1.3 × 10<sup>−5</sup> | 15–21 |
| Impaired Skeletal Muscle Contractility MP:0002841 | 4.7 × 10<sup>−3</sup> | 5.2 × 10<sup>−8</sup> | 10–17 |
| Impaired Skin Barrier Function MP:0002796 | 6.4 × 10<sup>−3</sup> | 1.2 × 10<sup>−8</sup> | 12–21 |
| Increased Susceptibility To Bacterial Infection MP:0002412 | 9.3 × 10<sup>−3</sup> | 2.2 × 10<sup>−7</sup> | 24–39 |
| Enlarged Spleen MP:0000691 | 1.1 × 10<sup>−2</sup> | 2.6 × 10<sup>−5</sup> | 55–72 |
| **Human_Phenotype_Ontology** |  |  |  |
| Palmoplantar keratoderma (HP:0000982) | ≥ 1 | 4.1 × 10<sup>−7</sup> | 0–28 |
| Skin ulcer (HP:0200042) | ≥ 1 | 2.5 × 10<sup>−6</sup> | 0–27 |
| Plantar hyperkeratosis (HP:0007556) | ≥ 1 | 2.1 × 10<sup>−5</sup> | 0–12 |
| Autosomal dominant inheritance (HP:0000006) | ≥ 1 | 1.2 × 10<sup>−3</sup> | 0–104 |
| Erythema (HP:0010783) | ≥ 1 | 1.2 × 10<sup>−3</sup> | 0–9 |
| Hypohidrosis (HP:0000966) | ≥ 1 | 1.2 × 10<sup>−3</sup> | 0–13 |
| Abnormal blistering of the skin (HP:0008066) | ≥ 1 | 1.4 × 10<sup>−3</sup> | 0–16 |
| Camptodactyly of finger (HP:0100490) | ≥ 1 | 1.5 × 10<sup>−3</sup> | 0–21 |
| Elbow flexion contracture (HP:0002987) | ≥ 1 | 2.2 × 10<sup>−3</sup> | 0–10 |
| Corneal erosion (HP:0200020) | ≥ 1 | 3.6 × 10<sup>−3</sup> | 0–10 |
| **Disease_Perturbations_from_GEO_up** |  |  |  |
| acute myocarditis DOID-3951 mouse GSE35182 sample 801 | 4.4 × 10<sup>−35</sup> | 3.3 × 10<sup>−53</sup> | 117–145 |
| Lung transplant rejection C0729958 mouse GSE3418 sample 83 | 1.1 × 10<sup>−34</sup> | 5.4 × 10<sup>−43</sup> | 79–95 |
| colitis DOID-0060180 mouse GSE34874 sample 800 | 2.2 × 10<sup>−27</sup> | 5.4 × 10<sup>−43</sup> | 81–103 |
| pulmonary tuberculosis DOID-2957 mouse GSE48027 sample 831 | 5.0 × 10<sup>−26</sup> | 3.1 × 10<sup>−43</sup> | 98–132 |
| cystic fibrosis DOID-1485 mouse GSE3100 sample 1057 | 1.1 × 10<sup>−25</sup> | 4.9 × 10<sup>−40</sup> | 77–101 |
| colon cancer DOID-219 mouse GSE63032 sample 661 | 1.2 × 10<sup>−25</sup> | 1.1 × 10<sup>−46</sup> | 85–104 |
| acute myocarditis DOID-3951 mouse GSE35182 sample 802 | 7.4 × 10<sup>−25</sup> | 5.2 × 10<sup>−45</sup> | 90–132 |
| Cystic Fibrosis C0010674 mouse GSE3100 sample 138 | 6.7 × 10<sup>−21</sup> | 3.9 × 10<sup>−33</sup> | 76–99 |
| basal cell carcinoma DOID-2513 mouse GSE58894 sample 730 | 9.2 × 10<sup>−19</sup> | 4.1 × 10<sup>−25</sup> | 58–66 |
| liver cirrhosis DOID-5082 human GSE50892 sample 1012 | 2.0 × 10<sup>−18</sup> | 2.7 × 10<sup>−30</sup> | 65–81 |
| **Disease_Perturbations_from_GEO_down** |  |  |  |
| psoriasis DOID-8893 mouse GSE27628 sample 823 | 7.3 × 10<sup>−22</sup> | 1.9 × 10<sup>−37</sup> | 70–87 |
| Neurological pain disorder C0423704 rat GSE15041 sample 356 | 6.9 × 10<sup>−16</sup> | 1.3 × 10<sup>−23</sup> | 71–77 |
| Barrett’s esophagus DOID-9206 human GSE34619 sample 453 | 2.5 × 10<sup>−11</sup> | 2.1 × 10<sup>−25</sup> | 50–75 |
| Barrett’s esophagus DOID-9206 human GSE34619 sample 596 | 1.4 × 10<sup>−10</sup> | 5.6 × 10<sup>−29</sup> | 48–78 |
| breast cancer DOID-1612 human GSE3744 sample 978 | 1.9 × 10<sup>−10</sup> | 1.0 × 10<sup>−23</sup> | 62–86 |
| Barrett’s esophagus DOID-9206 human GSE13083 sample 917 | 2.2 × 10<sup>−10</sup> | 6.5 × 10<sup>−21</sup> | 44–57 |
| basal cell carcinoma DOID-2513 mouse GSE58894 sample 730 | 3.9 × 10<sup>−10</sup> | 1.5 × 10<sup>−21</sup> | 47–74 |
| Breast Cancer C0006142 human GSE3744 sample 24 | 8.6 × 10<sup>−10</sup> | 1.4 × 10<sup>−21</sup> | 61–85 |
| pulmonary sarcoidosis DOID-13406 human GSE19976 sample 765 | 9.4 × 10<sup>−9</sup> | 1.2 × 10<sup>−21</sup> | 49–76 |
| Lung transplant rejection C0729958 mouse GSE3418 sample 83 | 2.2 × 10<sup>−8</sup> | 4.9 × 10<sup>−17</sup> | 45–58 |
| **Jensen_DISEASES_Curated_2025** |  |  |  |
| INTEGUMENTARY SYSTEM DISEASE | 7.2 × 10<sup>−4</sup> | 1.9 × 10<sup>−10</sup> | 31–48 |
| SKIN DISEASE | 2.1 × 10<sup>−3</sup> | 2.8 × 10<sup>−10</sup> | 25–42 |
| KERATOSIS | 2.0 × 10<sup>−2</sup> | 3.9 × 10<sup>−6</sup> | 9–14 |
| MUSCULOSKELETAL SYSTEM DISEASE | 2.7 × 10<sup>−2</sup> | 2.1 × 10<sup>−6</sup> | 44–69 |
| PALMOPLANTAR KERATOSIS | 3.4 × 10<sup>−2</sup> | 2.2 × 10<sup>−5</sup> | 8–12 |
| IMMUNE SYSTEM DISEASE | 3.5 × 10<sup>−2</sup> | 2.9 × 10<sup>−7</sup> | 22–41 |
| PRIMARY IMMUNODEFICIENCY DISEASE | ≥ 1 | 7.9 × 10<sup>−7</sup> | 0–37 |
| CORNEAL DISEASE | ≥ 1 | 3.3 × 10<sup>−5</sup> | 0–11 |
| AUTOIMMUNE DISEASE OF MUSCULOSKELETAL SYSTEM | ≥ 1 | 8.2 × 10<sup>−5</sup> | 0–25 |
| ICHTHYOSIS | ≥ 1 | 1.2 × 10<sup>−4</sup> | 0–7 |
| **DisGeNET** |  |  |  |
| Liver Cirrhosis, Experimental | 6.9 × 10<sup>−11</sup> | 1.3 × 10<sup>−19</sup> | 104–123 |
| Squamous cell carcinoma | 2.0 × 10<sup>−9</sup> | 5.3 × 10<sup>−19</sup> | 170–210 |
| Asthma | 2.8 × 10<sup>−9</sup> | 2.1 × 10<sup>−16</sup> | 126–160 |
| Neoplasm Metastasis | 4.9 × 10<sup>−9</sup> | 4.1 × 10<sup>−16</sup> | 298–352 |
| Psoriasis | 2.2 × 10<sup>−8</sup> | 4.2 × 10<sup>−12</sup> | 86–106 |
| Mammary Neoplasms | 2.8 × 10<sup>−8</sup> | 1.3 × 10<sup>−13</sup> | 186–238 |
| Degenerative polyarthritis | 3.6 × 10<sup>−8</sup> | 1.5 × 10<sup>−14</sup> | 94–128 |
| Adenocarcinoma | 1.0 × 10<sup>−7</sup> | 6.1 × 10<sup>−13</sup> | 152–182 |
| Pneumonia | 4.8 × 10<sup>−7</sup> | 1.5 × 10<sup>−17</sup> | 56–88 |
| Malignant neoplasm of breast | 9.4 × 10<sup>−7</sup> | 4.6 × 10<sup>−13</sup> | 342–422 |
| **GWAS_Catalog_2025** |  |  |  |
| Blood Protein Levels | 4.8 × 10<sup>−4</sup> | 1.9 × 10<sup>−8</sup> | 77–93 |
| Monocyte Percentage Of White Cells | ≥ 1 | 1.0 × 10<sup>−3</sup> | 0–46 |
| Eosinophil Percentage Of White Cells | ≥ 1 | 5.1 × 10<sup>−3</sup> | 0–43 |
| Allergic Disease (Asthma, Hay Fever And/Or Eczema) (Age Of Onset) | ≥ 1 | 1.1 × 10<sup>−2</sup> | 0–14 |
| Allergic Disease (Asthma, Hay Fever And/Or Eczema) (Multivariate Analysis) | ≥ 1 | 1.1 × 10<sup>−2</sup> | 0–14 |
| Asthma | ≥ 1 | 1.2 × 10<sup>−2</sup> | 0–71 |
| Hypertrophic Cardiomyopathy (Sarcomere Neg) | ≥ 1 | 1.4 × 10<sup>−2</sup> | 0–8 |
| Multiple System Atrophy | ≥ 1 | 3.3 × 10<sup>−2</sup> | 0–7 |
| Age-Related Macular Degeneration (Choroidal Neovascularisation) | ≥ 1 | 3.9 × 10<sup>−2</sup> | 0–4 |
| Aspartate Aminotransferase Levels | ≥ 1 | 3.9 × 10<sup>−2</sup> | 0–38 |
| **Drug_Perturbations_from_GEO_up** |  |  |  |
| soman 7305 rat GSE13428 sample 2637 | 1.2 × 10<sup>−30</sup> | 6.3 × 10<sup>−54</sup> | 97–124 |
| bleomycin DB00290 mouse GSE25640 sample 3121 | 2.4 × 10<sup>−30</sup> | 1.2 × 10<sup>−40</sup> | 89–111 |
| soman 7305 rat GSE13428 sample 2639 | 4.4 × 10<sup>−29</sup> | 7.9 × 10<sup>−45</sup> | 102–127 |
| ANGIOTENSIN II 172198 mouse GSE17901 sample 3678 | 1.7 × 10<sup>−28</sup> | 3.5 × 10<sup>−40</sup> | 83–100 |
| soman 7305 rat GSE13428 sample 2635 | 7.0 × 10<sup>−28</sup> | 1.1 × 10<sup>−51</sup> | 96–126 |
| soman 7305 rat GSE13428 sample 2640 | 4.1 × 10<sup>−27</sup> | 1.3 × 10<sup>−44</sup> | 100–125 |
| bleomycin DB00290 mouse GSE43695 sample 3104 | 1.1 × 10<sup>−25</sup> | 2.3 × 10<sup>−45</sup> | 73–97 |
| bleomycin DB00290 mouse GSE43695 sample 3106 | 4.4 × 10<sup>−24</sup> | 3.0 × 10<sup>−38</sup> | 82–96 |
| soman 7305 rat GSE13428 sample 2633 | 5.0 × 10<sup>−24</sup> | 3.7 × 10<sup>−39</sup> | 92–119 |
| phorbol ester 12-O-tetradecanoylphorbol-13-acetate (TPA) 27924 mouse GSE40849 sample 3257 | 3.5 × 10<sup>−21</sup> | 8.5 × 10<sup>−33</sup> | 59–73 |
| **Drug_Perturbations_from_GEO_down** |  |  |  |
| rosiglitazone 77999 mouse GSE11343 sample 2672 | 9.3 × 10<sup>−14</sup> | 1.4 × 10<sup>−23</sup> | 61–75 |
| formaldehyde 712 rat GSE7002 sample 3548 | 1.6 × 10<sup>−13</sup> | 2.2 × 10<sup>−22</sup> | 57–78 |
| sunitinib 5329102 rat GSE6929 sample 2491 | 4.6 × 10<sup>−13</sup> | 6.2 × 10<sup>−27</sup> | 69–92 |
| phorbol ester 12-O-tetradecanoylphorbol-13-acetate (TPA) 27924 mouse GSE40849 sample 3256 | 6.7 × 10<sup>−13</sup> | 4.5 × 10<sup>−26</sup> | 51–79 |
| dexamethasone DB01234 mouse GSE11056 sample 3358 | 2.6 × 10<sup>−12</sup> | 7.5 × 10<sup>−20</sup> | 52–70 |
| phorbol ester 12-O-tetradecanoylphorbol-13-acetate (TPA) 27924 mouse GSE40849 sample 3257 | 5.0 × 10<sup>−12</sup> | 1.7 × 10<sup>−17</sup> | 55–68 |
| pyrazinamide 1046 mouse GSE48027 sample 3326 | 4.4 × 10<sup>−10</sup> | 4.7 × 10<sup>−19</sup> | 47–69 |
| SODIUM ARSENITE 443495 mouse GSE11056 sample 3356 | 6.1 × 10<sup>−10</sup> | 1.1 × 10<sup>−21</sup> | 46–69 |
| etanercept DB00005 human GSE11903 sample 2608 | 1.8 × 10<sup>−9</sup> | 1.6 × 10<sup>−17</sup> | 43–60 |
| etanercept DB00005 human GSE41663 sample 2605 | 1.9 × 10<sup>−9</sup> | 4.2 × 10<sup>−17</sup> | 51–62 |

*S2* The same for the direction split, by time point and arm, over all six time points (36 760 rows), with a column marking the four that entered the comparison of Sec. 3.7 and the two that did not, whose rows are present in full.

*S3* The per-term classification behind the counts SAME only 74, OPPO only 87, BOTH 791.

*S4* Per library and time point, the number of terms, the number significant, and the number of submitted genes the library could match — the three quantities that make adjusted *P* and *k* incomparable between libraries.

*S5* The 23 gene lists behind every number stated in this paper, and no others, each labelled as primary, reported but not adopted, or superseded, with its gene count set beside the count printed here.

*S6* The long-form enrichment output including the member genes of every overlap.

## Declarations

### Ethics approval and consent to participate

Not applicable. This study analysed only data already archived in a public repository; no animal or human participants, data or tissue were involved in the work reported here.

### Consent for publication

Not applicable.

### Availability of data and materials

All data analysed in this study are publicly available from the Gene Expression Omnibus. The hippocampal bulk RNA-seq data are available under accession GSE292224, https://www.ncbi.nlm.nih.gov/geo/query/acc.cgi?acc=GSE292224; the hippocampal bulk ATAC-seq data under GSE292231, https://www.ncbi.nlm.nih.gov/geo/query/acc.cgi?acc=GSE292231 (both part of SuperSeries GSE292240); and the dorsal skin bulk RNA-seq data under GSE270713, https://www.ncbi.nlm.nih.gov/geo/query/acc.cgi?acc=GSE270713. No new data were generated. All derived results are in Additional file 1. The complete analysis code — download, the product-tensor runs, the chromatin-accessibility layer, enrichment, every table and figure in this paper, and a script that recomputes each published number and compares it against the value printed here — is available at https://github.com/tagtag/TDbasedUFE_Brain-Skin under the MIT licence. The underlying method is implemented in the Bioconductor packages TDbasedUFE and TDbasedUFEadv [7].

### Authors’ contributions

Y-HT conceived the study, curated the data, performed all analyses and wrote the manuscript. TT evaluated the results and reviewed and edited the manuscript. Both authors read and approved the final manuscript.

## Acknowledgements

Not applicable.

